# Spontaneous Pregravid Obesity Reshapes Fetal Immune Ontogeny in a Nonhuman Primate Model

**DOI:** 10.64898/2026.03.27.714880

**Authors:** Brianna M. Doratt, Sheridan Wagner, Uriel Avila, Travis Hodge, Lauren D. Martin, Oleg Varlamov, Ilhem Messaoudi

## Abstract

Pregravid obesity is associated with long⍰term immune alterations in the offspring; however, the mechanisms remain poorly defined. To address this gap, we investigated the impact of spontaneous pregravid obesity, independent of obesogenic diet, on fetal immune ontogeny in a rhesus macaque model. Using spectral flow cytometry, multiplex cytokine profiling, functional stimulation assays, and single⍰cell RNA sequencing, we profiled immune composition, function, transcriptional profiles, and intercellular communication in umbilical cord blood as well as fetal spleen and lung. Pregravid obesity was associated with altered fetal organ growth, elevated inflammatory mediators, altered frequencies of immune cell populations, and hyperresponsiveness to stimulation by splenic and lung leukocytes. Single⍰cell transcriptomic analyses revealed tissue⍰specific reprogramming of innate immune cells, including heightened inflammatory, migratory, and metabolic signatures with impaired antigen presentation. Moreover, there was evidence of impaired T cell differentiation, premature effector differentiation, and B cell dysfunction. Cell–cell communication analysis identified loss of tolerogenic signaling and enhanced proinflammatory pathways across spleen and lung myeloid cells. These findings demonstrate that spontaneous pregravid obesity fundamentally reshapes fetal circulating and tissue⍰resident immune cells, providing mechanistic insight into the increased susceptibility to infection, respiratory diseases, and immune dysregulation observed in offspring of mothers with obesity.

## Introduction

In the United States, over 41% and 12.1% of adult women were classified as obese (body mass index (BMI) > 30kg/m^2^) and severely obese (BMI > 40kg/m^2^) respectively in 2023(1-3). Pre-pregnancy (pregravid) obesity is associated with several adverse outcomes for pregnant women, including an increased risk of developing preeclampsia, gestational diabetes, hypertension, miscarriage and stillbirth(4-6). Moreover, pregravid obesity is a critical determinant of long-term offspring health, being associated with increased rates of allergies, asthma, necrotizing enterocolitis, sepsis, and severe respiratory syncytial virus (RSV) infection in offspring(7-19). Collectively, these outcomes suggest that pregravid obesity disrupts immune system development and maturation *in utero*.

Indeed, we have previously shown that umbilical cord blood (UCB) from neonates born to mothers with obesity generated blunted responses to in vitro stimulation with higher frequencies of memory T cells and a reduced frequencies of circulating monocytes(20-24). Data from rodent models of high fat or western style diet (HFD, WSD) induced obesity demonstrate increased susceptibility to bacterial challenge, hyper-reactive immune responses to infection, and increased IgE responses(17, 25, 26). Although rodent studies have provided important insights, their translational relevance is constrained by marked differences between rodent and human immune ontogeny. Clinical studies also face significant limitations in evaluating tissue⍰resident leukocyte populations and are frequently confounded by environmental and lifestyle variables that are difficult to control.

Nonhuman primate (NHP) models provide a unique opportunity to uncover mechanistic insights in a genetically outbred animal model while controlling for confounders that challenge clinical studies. Leveraging this model, we have shown that pregravid obesity impacts the fetal bone marrow (FBM) milieu and hematopoietic stem and progenitor cell (HSPC) differentiation trajectories in both diet-induced and spontaneous NHP models of pregravid obesity (27-29). Other studies using this model have reported disruptions in offspring metabolic health and altered expression of genes critical for antigen presentation, complement and coagulation, leukocyte migration, and B⍰cell receptor signaling(23, 30-35). However, current NHP models of pregravid obesity rely almost exclusivly on high-fat, calorie-dense WSD, which fail to recapitulate the full spectrum of human obesity etiologies, including physical inactivity, environmental exposure, and genetic predisposition influencing basal metabolism and appetite(36-39).

Therefore, this study examined alterations in circulating and tissue⍰resident leukocytes in gestational day (GD) 130 fetuses derived from lean or obese rhesus macaques fed a regular chow diet. Rhesus macaque gestation closely parallels human pregnancy, with GD130 corresponding to the mid–third trimester(40, 41). We profiled umbilical cord blood mononuclear cells and leukocytes isolated from the spleen and lung. Immune phenotyping and functional assessment were performed using spectral flow cytometry, in vitro stimulation assays, and single⍰cell RNA sequencing (scRNA⍰seq). We report that pregravid obesity is associated with altered seeding frequencies of tissue⍰resident immune cells and a shift toward a more proinflammatory immune profile.

## Methods

### Cohort description

This study was approved by the Oregon National Primate Research Center (ONPRC) Institutional Animal Care and Use Committee and conforms to current Office of Laboratory Animal Welfare regulations (assurance number A3304-01). Adult female rhesus macaques were pair-housed, with the cage size adjusted per USDA Cage Size Guide, eighth Edition. Animals were maintained on a monkey chow diet consisting of two daily meals of fiber-balanced monkey diet (15% calories from fat, 27% from protein and 59% from carbohydrates; no. 5052; Lab Diet, St. Louis, MO), supplemented with fruits and vegetables. Six lean and seven spontaneously obese adult female rhesus macaques were selected from the ONPRC colony based on the body condition score (BCS). BCS is a semiquantitative method of assessing body fat and muscle, on a scale of 1-5, by palpation of key anatomic features including the hip bones, facial bones, spinous processes and ribs as well as muscle mass present over the hip bones and prominence of the ischial callosities(42). Obese animals were defined as having a BCS ≥4, with lean animals having a BCS of 2-3. All animals in this study were managed according to the ONPRC animal care program, the AAALAC International, and guidelines set forth by the United States Department of Agriculture.

### Time mated breeding

Indian origin rhesus macaques (*Macaca mulatta*) were assigned to the study from the ONPRC time-mated breeding (TMB) colony. Adult females were monitored for menstruation and non-sedated daily blood draws were initiated at the mid-follicular phase to assess serum estradiol and progesterone levels in order to determine optimal male pairing. Beginning 8 days after the onset of menses, the female was pair-housed with a mature male of proven fertility for 7-10 consecutive days. Pair compatibility was monitored during co-housing. At approximately 30Ldays post-mating, females were sedated with ketamine hydrochloride (8-10 mg/kg). Three-dimensional/four-dimensional (3D/4D) ultrasonography was performed using a GE Voluson E8 system (GE Healthcare, Chicago, IL) equipped with an RNA5-9-D 4D micro-convex transducer to visualize the gestational sac, when present, and estimate fetal gestational age based on crown–rump length measurements(43).

### Metabolic and hormone assays

Dual-energy X-ray absorptiometry (DEXA), intravenous glucose tolerance tests (GTTs), and measurements of fasting glucose and insulin were conducted as previously described(44-47). For DEXA assessments, monkeys were sedated with ketamine and positioned supine on a Hologic Discovery DEXA scanner (Hologic Inc., Bedford, MA, USA). Two to three scans were acquired per animal, and mean values were used for analysis. For GTTs, animals were fasted overnight and sedated with Telazol (3–5 mg/kg) prior to administration of a glucose bolus (50% dextrose) at 0.6 g/kg via the saphenous vein. Blood samples were collected immediately before glucose infusion and then 1, 3, 5, 10, 20, 40, and 60 minutes post-injection by venipuncture. Blood glucose levels were measured using a OneTouch Ultra Blood Glucose Monitor (LifeScan). Insulin was analyzed in the ONPRC Endocrine Technologies Core by immunoassay on a Roche cobas e411 chemiluminescence-based automated immunoassay platform (Roche Diagnostics, Indianapolis, IN). The assay range was 0.2-1000 μIU/mL. Intra-assay CV was 1.1%. Inter-assay CV for the insulin assay is 3.7%. Insulin resistance was estimated using the Homeostatic Model Assessment of Insulin Resistance (HOMA-IR), calculated as fasting serum insulin (μU/mL) × fasting plasma glucose (mg/dL) / 405.

### Tissue processing

Blood samples were collected in EDTA vacutainer tubes. Plasma and PBMC/UCBMC were isolated after centrifugation over lymphocyte separation media (STEMCELL Technologies). Splenocytes were isolated by mechanical disruption and filtered through a 70Lμm cell strainer followed by lysis to remove red blood cells. Lung tissue was digested for 1hr at 37C using 3mg/mL collagenase II, 0.2U/mL elastase, 1mg/mL DNAse, 0.3mg/mL hyaluronidase, 0.221 mg/mL CaCl2 in RPMI. The cell suspension was filtered through a 100 μm cell strainer and layered over a 60%/30% Percoll gradient, centrifuged, and mononuclear cells were collected from the 30%/60% interface. Plasma was stored at -80^0^C and PBMC,UCMBC, splenocytes, and lung leukocytes were cryo-preserved using 10% DMSO/FBS in a cryogenic unit until analysis.

### Spectral flow cytometry

UCMBC, splenocytes, and lung leukocytes were stained with a surface antibody cocktail in the presence of 50 µl Brilliant Stain Buffer, 5 µl TruStain rhesus FcX, and 5 µl True-Stain Monocyte Blocker for 30 minutes at 4°C in the dark using the following panel: CD163 (PE), CD14 (BV711), CD11c (BV421), CD123 (PE-CF594), HLA-DR (APC-Cy7), CD16 (BUV395), CD3 (BV650), CD8A (CFLOUR B548), CD4 (APC), CCR7 (BV785), CD28 (PE-CY7), CD95 ( BUV615), CD20 (PerCP-Cy5.5), IgD (Biotin, with Streptavidin BV510 as a secondary), CD27 (CFLOUR YG584). Cells were washed twice before fixation with Cytek FoxP3 Fix/Perm buffers for two hours at 4°C. Cells were intranuclearly stained with Ki-67 (FITC) overnight at 4°C. Cells were washed and samples were analyzed using the SpectroFlo Software (v3.0) and a Cytek Aurora spectral flow cytometer equipped with five lasers. Spectral unmixing was performed using single-stained reference controls (beads or cells), with autofluorescence extraction enabled. Unmixed FCS files were analyzed using FlowJo (v10.10.0).

### Luminex analysis and ligand stimulation

Pasma samples were assayed using the Nonhuman Primate (NHP) XL Cytokine Luminex Performance Premixed Kit (R&D Systems, FCSTM21⍰36) following the manufacturer’s instructions. The panel quantified the following 36 analytes: BDNF, CXCL13, CCL11, CCL2, CD40L, CCL20, CXCL10, CXCL11, FGFb, G-CSF, GM-CSF, IFNb, Granzyme B, IFNa, IFNg, IL1b, IL-10, IL-12p70, IL-13, IL-15, IL-17A, IL-5, IL-2, IL-21, IL-4, PDL1, IL-6, IL-7, IL-8, PDGF-AA, PDGF-BB, CXCL2, CCL5, TGFa, TNFa, and VEGF. UCMBC, splenocytes, and lung leukocytes were thawed and rested for 2 hours before stimulation with various ligand cocktails for 24 hours at 37°C. The PMA/IONO cocktail consisted of 0.05 μg/mL of PMA and 1 μg/mL of Ionomycin. The bacterial ligand cocktail consisted of 1 μg/mL Pam3CSK4, 0.5 μg/mL of FSL-1, and 0.5 μg/mL LPS. The viral ligand cocktail consisted of 1 μg/mL ssRNA40, 5 μg/mL Imiquimod, and 1 μg/mL ODN2216. All stimulation ligands were purchased from Invitrogen. Supernatants from the stimulations were collected and run on a custom NHP Millipore Luminex kit using manufacturer’s protocol to assess the following 28 analytes: BDNF, CXCL13, CCL11/Eotaxin, CCL2, CD40L, CCL20, CXCL10, CXCL11, FGFbasic, G-CSF, GM-CSF, IFN-b, GZMB, IFN-a, IFN-g, IL-1b, IL-10, IL-12, IL-13, PD-L1, IL-6, IL-8, PDGF-AA, PDGF-BB, CXCL2, CCL5, TNF-a, and VEGF.

### 3′multiplexed single-cell RNA sequencing

Freshly thawed cells were incubated with Fc blocker in PBS with 1% BSA for 10 min at 4^0^C and then surface-stained with CD45-FITC and a unique hashtag oligo (HTO) for 30 min in the dark. Samples were then washed in PBS with 0.04% BSA, stained with PI for live dead discrimination, and live CD45+ cells were then sorted on a Sony SH800 into RPMI (supplemented with 30% FBS). Sorted live leukocytes were counted in triplicates on a TC20 Automated Cell Counter (BioRad), washed, and resuspended in PBS with 0.04% BSA in a final concentration of 1600 cells/uL. Single cell suspensions were then immediately loaded on the 10X Genomics Chromium X with a loading target of 30,000 cells. Libraries were generated using the V3.1 chemistry (for gene expression) and Single Cell 3L Feature Barcode Library Kit per the manufacturer’s instructions. Libraries were sequenced on Illumina NovaSeq X with a sequencing target of 30,000 gene expression reads and 5,000 HTO reads per target cell.

### Single-cell RNA-Seq data analysis

For 3′ gene expression, raw reads were aligned and quantified using Cell Ranger (version 7.2.0, 10X Genomics) against the Mmul_8.0.1 reference genome using the *count* option. HTO doublets were then removed in Seurat using the *HTODemux* function. Droplets with ambient RNA (< 400 detected genes), cell doublets (> 4000 detected genes), and dying cells (>20% mitochondrial gene expression) were excluded during initial QC. Batch correction was performed using Harmony package. Data normalization and variance stabilization were performed on the integrated object using the *NormalizeData* and *ScaleData* functions in Seurat. Dimension reduction was performed using *RunPCA* function and clusters were visualized using Seurat’s *RunUMAP*.

Cell types were assigned to individual clusters using canonical gene markers from the *FindAllMarkers* function with a log_2_ fold change cutoff of at least 0.4 and an FDR<0.05 (Supp Table 1). Differential gene expression analysis was performed using the Wilcoxon rank-sum test using default settings in Seurat. Only statistically significant genes (log_2_(fold change) cutoff ≥ 0.4; adjusted p-value ≤ 0.05) were included in downstream analyses (Supp Table 2). Module scores for specific gene sets were incorporated cluster-wise using *AddModuleScores* function.

The R CellChat(48) package was employed to infer probable intercellular communication networks, utilizing the CellChatDB.human database. Data was preprocessed with the function identifyOverExpressedGenes and identifyOverExpressedInteractions and communication probabilities between clusters were determined with computeCommunProb (truncatedMean, trim=0.1, interaction.range=250, contact.range=100). Communications were filtered (filterCommunication) to a minimum number of 10 cell and signaling pathway probabilities were calculated (computeCommunProbPathway).

### Statistical Analysis

GraphPad Prism 10 was used for data analysis. Normality testing was performed using the Shapiro-Wilk test and outliers were assessed using the ROUT test with Q=1%. Two group comparisons were compared using a Welch’s t-test, while four-way comparisons of stimulated and non-stimulated conditions were assessed using a one-way ANOVA with Bonferroni correction. For all figure panels error bars indicate the standard error of the mean and p-values may be represented by symbols defined as follows: # = p-value <0.1, *= p-value <0.05, ** = p-value <0.01, *** = p-value <0.001, **** = p-value <0.0001.

## Results

### Maternal Body Composition, Metabolism and Cytokine Production

Obese and lean females were selected from the NHP colony, and DEXA analyses were performed at baseline to assess body composition, including lean and fat mass (Fig. 1A). Lean dams had a higher percentage of lean mass compared to fat mass, while those with pregravid obesity had a significantly higher percentage of fat mass, especially in the lower-body (gynoid) and upper-body (android) regions (Fig. 1A). Maternal metabolic state was evaluated at gestational days (GD) 0 (baseline), 30, 90, and 130. Obese dams had consistently higher weight throughout gestation. As previously reported in humans (49, 50), lean dams gained significantly more weight during gestation than obese dams, beginning at gestational day (GD) 90, whereas the body weight of obese dams remained relatively stable across gestation (Fig. 1B).

**Figure 1:**
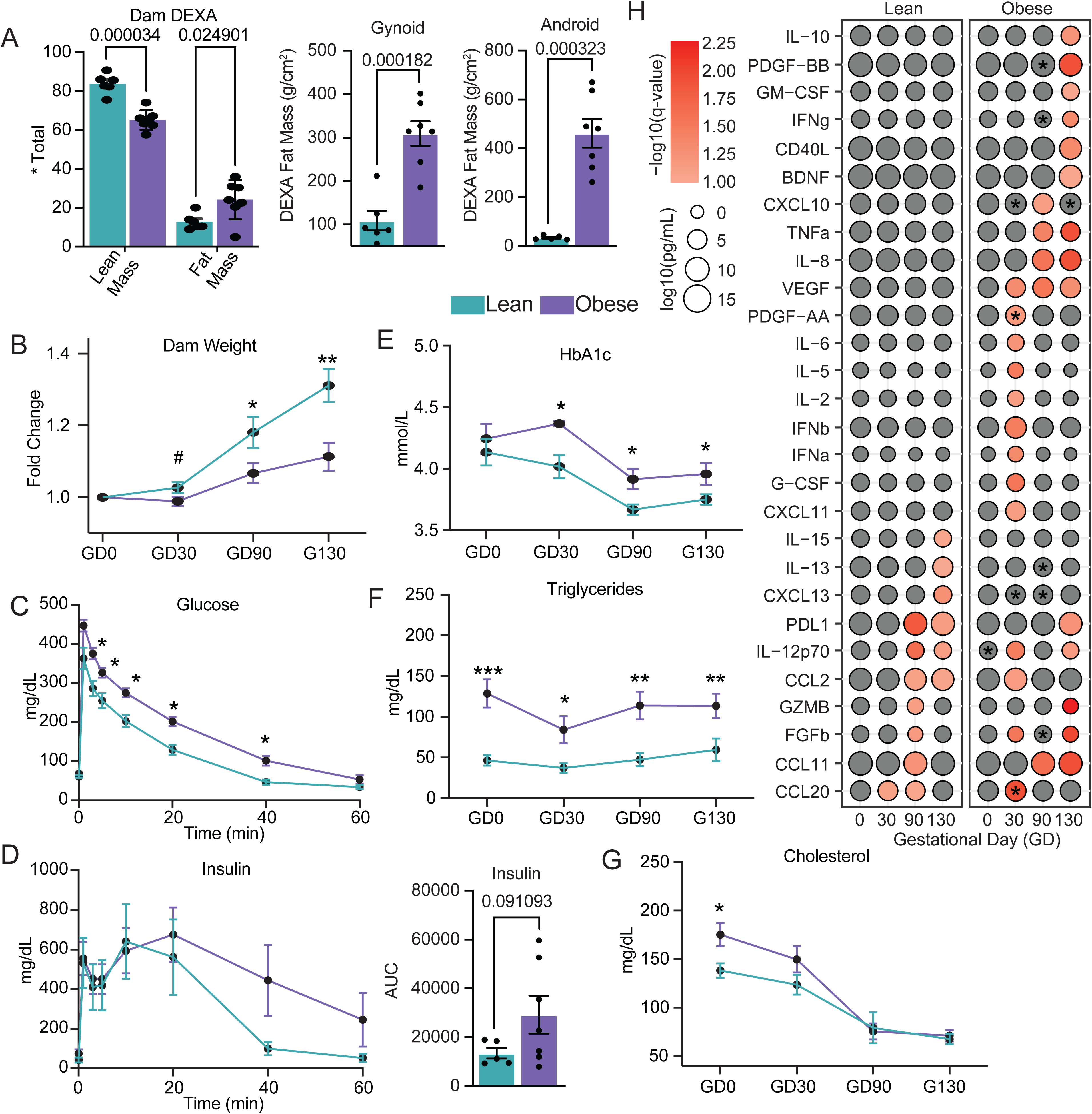
Metabolic and inflammatory characterization of dams with spontaneous obesity. A) Body composition analysis by DEXA showing percent of total mass for lean mass and fat mass (lest), total mass of gynoid fat mass (middle) and android fat mass (right). B) Longitudinal measurements of dam weight throughout gestation as a fold change from baseline, C-D) Glucose tolerance test (GTT) performed pre-pregnancy, showing glucose and insulin levels. E-G) Longitudinal measurements of HbA1c, circulating levels of triglycerides and cholesterol throughout gestation. H) Bubble plot of cytokine production throughout gestation in dam plasma measured by Luminex. Bubble size represents the concentration in log10(pg/mL), while bubble color reflects the -log10(qvalue) comparing the timepoint to GD0 for the corresponding group. *p < 0.05 by direct comparison of lean and obese samples at the indicated timepoint.

Glucose tolerance testing (GTT) revealed significantly greater glucose intolerance in obese dams, indicated by slower glucose clearance rates (Fig. 1C), with a trending increase in insulin area under the curve (AUC) during the GTT (Fig. 1D). However, fasting glucose levels were significantly decreased by AUC analysis in obese dams, while AUC of fasting insulin levels showed a significantly increased across gestation (Supp Fig 1A,B). This pattern is consistent with a state of chronic hyperinsulinemia and insulin resistance, reinforcing the metabolic burden carried into late gestation. AUC hemoglobin A1c (HbA1C) levels were significantly elevated throughout pregnancy with pregravid obesity, albeit without reaching the threshold indicative of diabetes (HbA1c>5%)(51) (Fig. 1E). Additionally, spontaneous pregravid obesity was associated with significantly increased triglyceride levels throughout gestation, while cholesterol levels were significantly different only at baseline (Fig. 1F,G).

We next determined the impact of pregravid obesity on circulating immune mediators at GD 0, 30, 90, and 130. Relative to baseline conditions, lean dams showed decreased levels of CCL20 at GD30, CCL11, FGFb, GZMB, CCL2 and IL-12p70 at GD90, and CXCL13, IL-13 and IL-15 at GD130 (Fig. 1H). Only levels of PDL1 were significantly increased at GD90 through GD130 in lean dams (Fig. 1H). Alternatively, in obese dams, there was an increase in the circulating levels of CCL2, CCL20, CXCL11, G-CSF, IFNα, IFNβ, IL-2, IL-5, IL-6, and VEGF at GD30, and PDL1 at GD130 (Fig. 1H). Conversely, the levels of PDGF-AA and FGFb were decreased at GD30, CCL11, IL-8, CXCL10, and TNFα levels were decreased at GD90, and BDNF, CD40L, GM-CSF, GZMB, IFNγ, IL-10, and PDGF-BB were decreased at GD130 in the obese dams (Fig. 1H). Next, we directly compared the analyte levels between the lean and obese dams at each gestational timepoint. At baseline (GD0), the levels of IL-12p70 were significantly decreased in the obese group compared to lean controls (Fig. 1H). At GD30, the levels of PDGF-AA were also decreased while the levels of CCL20, CXCL10 and CXCL13 were increased in obese dams (Fig. 1H). Finally, at GD90, levels of IFNγ and IL-13 were decreased, while levels of FGFb and PDGF-BB were increased in obese dams (Fig. 1H). Finally, looking at an area under the curve (AUC) analysis, levels of CXCL13, CXCL11, IL-12, IL-2, IL-6, CXCL10, PDGF-BB, IL-7, and CCL20 were higher while levels of IL-10, TGFα, IFNγ, IL-13, and BDNF were decreased throughout gestation with pregravid obesity (Supp Fig 1C).

### Fetal Anthropometry, Cytokine Profiles, and Cellular Composition of UCB

Placental weight and overall fetal body weight were significantly increased with pregravid obesity, with no change in crown-rump length (Fig. 2A). Interestingly among the fetal organs, pancreas and adrenal gland weights were significantly decreased, while the heart and lung weights were significantly increased with pregravid obesity (Fig. 2A). No changes in the weights of the pituitary gland, spleen, kidneys, liver, thymus, and brain were noted (Supp Table 3). These observations suggest that pregravid obesity appears to enhance development of cardiovascular and respiratory tissues while impairing endocrine organ development.

**Figure 2:**
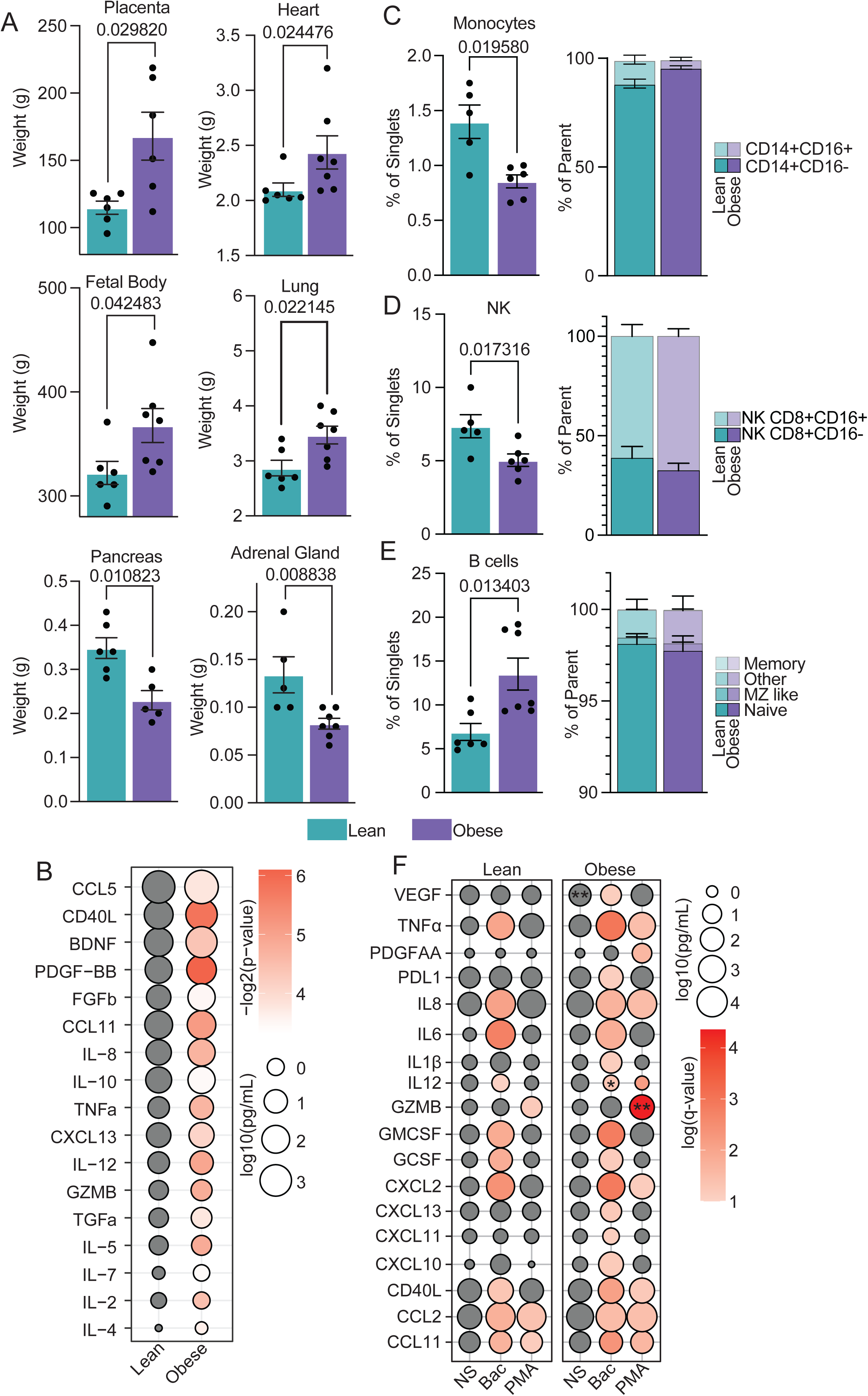
Pregravid obesity alters fetal cytokines and blood immune cell composition. A) Placental, fetal body, and select fetal organ weights. B) Bubble plot of cytokine concentrations in fetal plasma measured by Luminex. Bubble size represents cytokine abundance log10(pg/mL), while bubble color reflects the -log(qvalue) from multiple t-test comparisons of lean vs obese. C-E) Bar plots showing the frequency of the indicated cell types in each group using spectral flow cytometry. Gating strategy is shown in Supplementary Fig. 2A. F) Bubble plot of cytokine production following ligand cocktail stimulation (Bac, bacterial, PMA). Bubble size represents cytokine concentration log10(pg/mL), while bubble color reflects the q-value from a two-way ANOVA comparing stimulated conditions to the non-stimulated (NS) condition within each group. *p < 0.05, **p < 0.01 for comparisons of cytokine production after stimulation in obese relative to lean groups.

We also measured immune mediator levels in UCB plasma. Levels of CCL5, CD40L, BDNF, PDGF-BB, FGFb, CCL11, IL-8, IL-10, TNFα, CXCL13, IL-12, GZMB, TGFα, IL-5, IL-7, IL-2, and IL-4 were elevated with pregravid obesity (Fig. 2B). Given the altered cytokine and chemokine profiles observed in UCB, we assessed changes in immune cell composition in UCBMC using spectral flow cytometry (Supp Fig 2A). The frequency of total monocytes decreased, with a relative increase in classical CD14+CD16- monocytes (Fig. 2C). Similarly, the frequency of natural killer (NK) cells was significantly reduced with pregravid obesity, with a modest increase in the cytotoxic CD16+ subset (Fig. 2D). Conversely, the frequency of total B cells was significantly increased with pregravid obesity, with no changes observed among naïve/memory subsets (Fig. 2E). Finally, no differences were detected in the frequency of dendritic cells (DC), including the myeloid (mDC) and plasmacytoid (pDC) subsets, or in any T cell subset (Supp Fig 2B-C).

To assess the impact of pregravid obesity on the functional capacity of UCBMC, total UCBMC were stimulated with bacterial ligands or PMA/ionomycin, and immune mediators were measured by Luminex. Bacterial ligand stimulation elicited a robust response in the obese group, with elevated levels of VEGF, PDL1, IL1B, CXCL10, and CXCL11 uniquely in the obese group (Fig. 2F). In contrast, PMA stimulation induced a more limited response, with only CCL11, CCL2, and GZMB increased in both groups with GZMB induction significantly higher in the obese group (Fig. 2F). Additionally, CD40L, CXCL2, IL8, PDGF-AA, and TNFα were uniquely elevated in the obese group following PMA stimulation (Fig. 2F). Interestingly, although IL-12 was significantly induced by bacterial stimulation in the lean group, levels decreased in the obese group compared to baseline under both bacterial and PMA stimulation (Fig. 2F). Finally, VEGF levels were significantly higher in the resting NS supernatant from the obese group (Fig. 2E).

### Impact of Pregravid obesity on Gene Expression in UCB Innate Immune Cells

To identify molecular mechanisms underlying functional differences observed between the obese and lean groups, we performed scRNAseq. We identified 12 clusters based on canonical marker gene expression (Fig. 3A-C and Supp. Fig 2D). T cell clusters were identified by high *CD3* expression and comprised a CD8 T cell cluster (*CD3+CD8+*), a Treg cluster (*IL2RA+*), two CD4 T cells clusters (*CD3+CD8-)* distinguished by differential expression of *IL7R* and *CD28,* and a cluster of proliferating T cells (*MKI67+*). The NK cell compartment was identified as *CD3-CD8+NKG2D+GZMB+* cells (Fig. 3A,B). A population of NKT cells was identified based on co-expression of *CD3* and the transcription factor *ZBTB16* (Fig. 3A,B). Two B cell clusters (*MS4A1+*) with differential expression of *IL7R* were also identified (Fig. 3A,B). The myeloid compartment was comprised of the monocyte cluster (*MAMU-DRA+CD14+)* and the DC cluster (*MAMU-DRA+CD1C+*) (Fig. 3A,B). UCBMC also contained a small population of hematopoietic stem and progenitor cells (HSPC, *CD34+*) (Fig. 3A,B).

**Figure 3:**
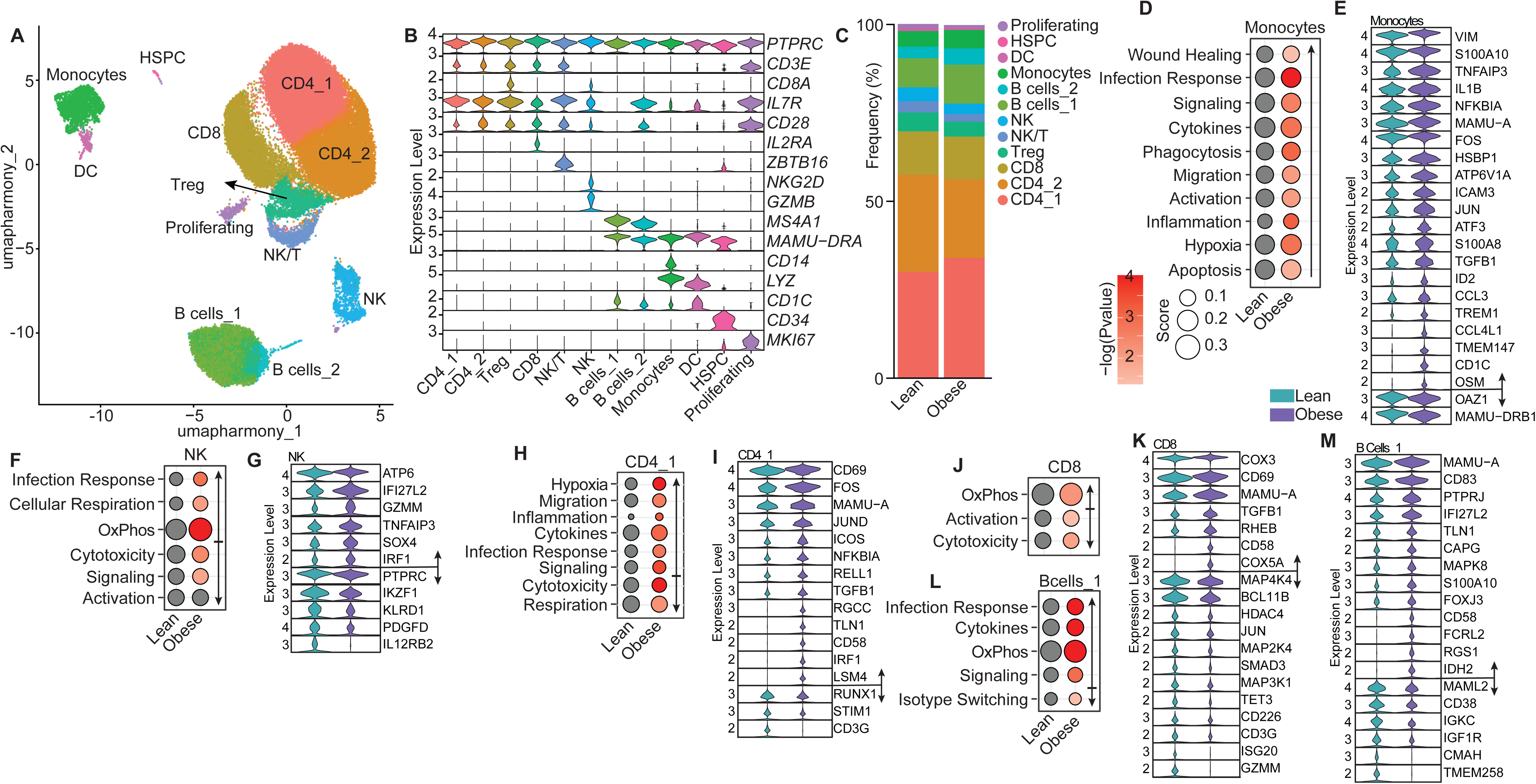
Impact of pregravid obesity on gene expression in umbilical cord blood. A) UMAP visualization of scRNAseq analysis of UCBMCs, with colors indicating distinct cell clusters. B) Violin plots of marker genes used to define each immune cell cluster. C) Stacked bar plot showing relative cluster frequencies across groups. D, F, H, J, L) Bubble plots depicting module scores for the indicated clusters. Bubble size represents the average module score, while bubble color reflects the p-value from multiple t-test comparisons. E, G, I, K, M) Violin plots showing expression of the indicated differentially expressed genes (DEGs) within the indicated clusters.

Although the frequency of cell clusters was not affected by pregravid obesity (Fig. 3C), module scoring and differential gene expression (DEG) analysis predicted transcriptional shifts associated with pregravid obesity (Fig. 3D-M). Scores for wound healing, infection response, signaling, phagocytosis, migration, hypoxia, and apoptosis were increased in monocytes with pregravid obesity (Fig. 3D). This was accompanied by upregulation of AP-1 transcription factors (*FOS, JUN*), NF-κB signaling (*NFKBIA, TNFAIP3*), migration (*ICAM3, VIM*), stress (*HSBP1*), inflammatory mediators (*S100A8, S100A10, IL1B, CCL3, CCL4L1*), and antigen presentation receptors (*TREM1, CD1C*) (Fig. 3E). Interestingly, HLA class I genes (*MAMU-A*) were increased while HLA class II genes (*MAMU-DRB1*) were decreased (Fig. 3E). Both NK and NKT cell clusters showed increased scores for infection response, respiration, and OxPhos modules, accompanied by decreased scores for cytotoxicity and signaling (Fig. 3F, Supp. Fig 2E). Despite the overall decrease in the NK cell cytotoxicity module, we observed upregulation of the *GZMM* gene (Fig. 3G). In addition, expression of genes associated with interferon (IFN) responses (*IFI27L2, IRF1*) were increased while genes critical for NK cell differentiation and education (*IKZF1, KLRD1*) were decreased (Fig. 3G).

**T**he CD4_1 cluster exhibited more pronounced transcriptional changes in response to pregravid obesity than the CD4_2 cluster. Scores associated with hypoxia, inflammation, migration and signaling modules were increased in the CD4_1 cluster while respiration module was decreased (Fig. 3H). Indeed, the early activation marker *CD69*, T cell co-stimulatory molecules (*ICOS, CD58*), as well as genes involved in cell cycle and protein synthesis (*LSM4, RGCC*) were upregulated with pregravid obesity (Fig. 3I). Interestingly, pregravid obesity was associated with reduced expression of the canonical T cell genes, including *CD3G*, a subunit of the T cell receptor (TCR), and *RUNX1*, a transcription factor essential for T cell maturation (Fig. 3I). CD8 T cells from the obese group exhibited increased scores for OxPhos module and decreased scores associated with activation and cytotoxicity (Fig. 3J). CD8 T cells also showed upregulation of the early activation marker *CD69* and the co-stimulatory molecule *CD58*, along with downregulation of the TCR subunit *CD3G*, with pregravid obesity (Fig. 3K). In addition, mitochondrial electron transport chain genes (*COX3, COX5A*) and *TGFB1*, which inhibit cytotoxic of T cells functions (52, 53) were upregulated (Fig. 3K). Expression of *GZMM*, MAPK factors (*MAP4K4*, *MAP2K4*), epigenetic regulators (*HDAC4*, *TET3*), co-stimulatory molecule CD266 (*DNAM-1*), and transcription factor *BCL11B* were decreased with pregravid obesity (Fig. 3K).

The Bcell_1 cluster exhibited increased scores for infection response, signaling, and OxPhos, with decreased scores for isotype class switching module (Fig. 3L). With pregravid obesity, fetal B cells showed a decrease in the expression of *IGF1R*, a tyrosine kinase essential for cell survival and proliferation, and *IGKC*, the most abundant immunoglobulin light chain (Fig. 3M). Additionally, we observed upregulation of the activation marker *CD83*, genes involved in cell migration (*CAPG*, *TLN1*), the stress-induced *MAPK8*, the mitochondrial enzyme *IDH2*, *RGS1*, a signal regulator linked to decreased chemokine signaling in B cells, *FCRL2*, an inhibitory receptor typically expressed on memory B cells, and *PTPRJ*, a tyrosine phosphatase associated with dampened BCR activation (Fig. 3M)(54-58).

### CellChat Analysis on the scRNAseq UCBMC Dataset

We next assessed potential alterations in the signaling pathways within UCBMC using CellChat. We observed no change in the overall count of potential interactions with pregravid obesity however the overall interaction strength was decreased (Fig 4A). When examining cluster-to-cluster communication, interaction counts were decreased for NK/T cells, DCs and Proliferating cells, while interaction counts were increased from HSPC to all other clusters, Monocytes to T and NK cell clusters, and CD4 Tcells to multiple clusters (Fig 4B). Similarly, cluster-by-cluster interaction strength was largely decreased, except for CD4_1 signaling to CD4_1 and CD4_2 (Fig 4B). Next, we quantified the relative information flow, an aggregate measure of both the number and strength of the signals for each pathway in each group (Fig. 4C). Several signaling pathways were predicted to be stronger in the lean group, including PDGF, CD39, CD48, and CD40 (Fig. 4C). Pathways predicted to be stronger in the obese group included CD96, SEMA7, and SLURP (Fig. 4C). Most of the changes in signaling pathways were observed in the monocyte, DC, and HSPC populations (Fig.4D). CD23, a low-affinity receptor for IgE, showed increased signaling to monocytes and decreased signaling to DC with pregravid obesity (Fig.4D,E)(59). Signling with PCDH, a cadherin that modulates actin cytoskeleton dynamics, was attenuated with pregravid obesity and shifted to the HSPC cluster with pregravid obesity (Fig.4D,F). CD39 signaling, a key ectonucleotidase that initiates the breakdown of pro-inflammatory extracellular ATP, was largely lost in monocyte and DC clusters with pregravid obesity (Fig.4D,G)(60). Signaling of PECAM1, an immunoregulatory receptor that facilitates leukocyte migration, to monocytes and the proliferating cluster from T cell clusters increased with pregravid obesity (Fig.4D,H).

**Figure 4:**
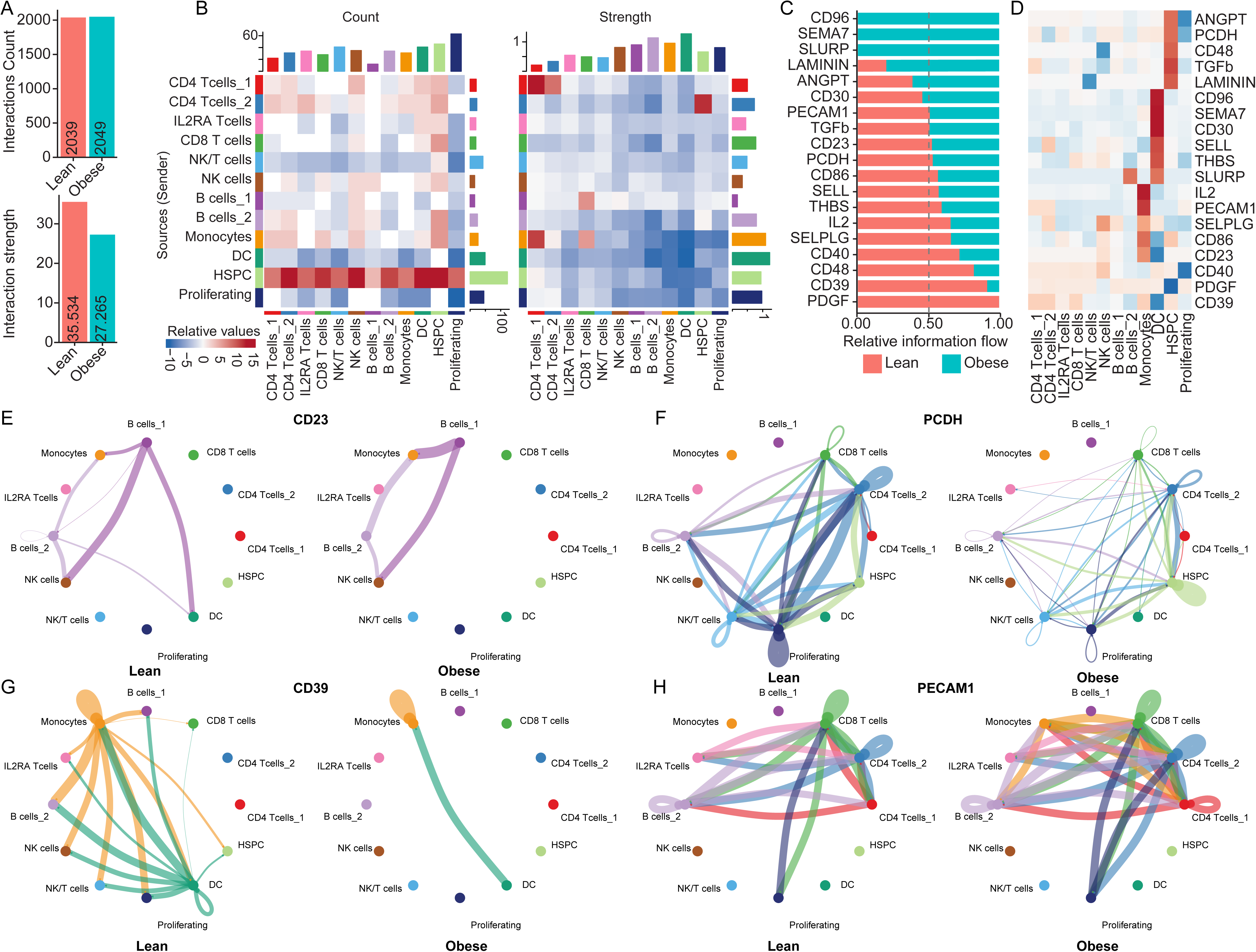
Pregravid obesity alters cell-cell communications in umbilical cord blood. A) Bar plot showing the inferred number and strength of cell–cell interactions using CellChat analysis on the scRNAseq UCBMC data. B) Heatmap depicting relative interaction counts and interaction strengths between clusters. Blue indicates reduced interactions in the obese group compared with the lean group. Bars along the top and right margins indicate cumulative signaling interactions for each column and row, respectively. C) Bar plot showing relative information flow, defined as the aggregate probability of communication, for each signaling pathway. D) Heatmap illustrating differential cluster-to-pathway interactions. Red indicates increased interaction potential in the obese group relative to the lean group. E–H) Circle plots showing cluster-to-cluster communication for the indicated signaling pathways. Lean samples are shown on the left and obese samples on the right. Line color denotes the source cell population, and line thickness reflects the relative interaction count.

### Impact of Pregravid obesity on Spleen Immune Landscape

Next, we assessed the impact of pregravid obesity on the immune landscape of the spleen. Using spectral flow cytometry, we observed an increase in the frequency of splenic macrophages, pDC, and B cells with no changes in the frequencies of mDC or NK cells with pregravid obesity (Fig 5A-C, Supp Fig 3A-C). We noted no change in the frequency of CD8 T cells but a modest reduction in the frequency of CD4 T cells with pregravid obesity (Fig 5D, Supp Fig 3D). Within the CD4 T cells, we observed an increase in the frequency of terminal effector memory (TEM) cells at the expense of naïve cells and no change in the frequency of effector memory (EM) or central memory (CM) cells (Fig. 5D).

**Figure 5:**
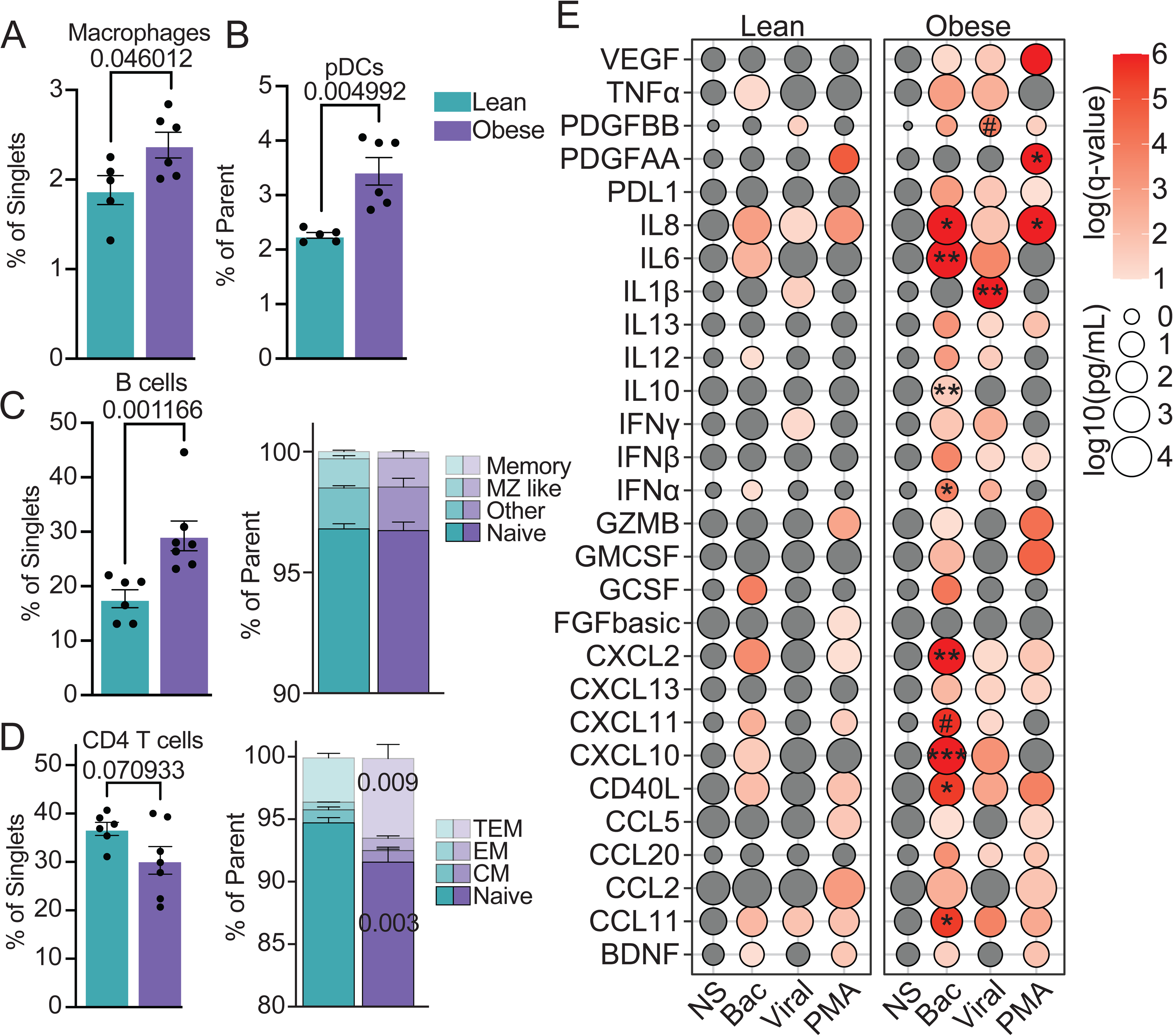
Impact of pregravid obesity on spleen immune landscape. A–D) Bar plots showing the frequency of the indicated cell types in each group. The gating strategy, using spectral flow cytometry, is shown in Supplementary Fig. 3A. E) Bubble plot of cytokine production following ligand cocktail stimulation. Bubble size represents cytokine concentration log10(pg/mL), while bubble color reflects the p-value from a two-way ANOVA comparing stimulated conditions to the non-stimulated (NS) condition within each group. #p < 0.1, *p < 0.05, **p < 0.01 for comparisons of cytokine production after stimulation in obese relative to lean groups.

We next assessed functional changes of splenic leukocytes with pregravid obesity using in vitro stimulation (Fig 5E). While stimulation with bacterial, viral, and PMA cocktails elicited responses in both groups, the obese group generated a heightened response with the production of multiple uniquely produced cytokines that were not induced in lean controls (Fig 5E). Specifically, CCL2, CCL20, CCL5, CXCL13, GM-CSF, GZMB, IFNβ, IFNγ, IL-10, IL-13, PD-L1, PDGF-BB, and VEGF were significantly increased following bacterial stimulation relative to the NS control only in splenocytes from the obese group (Fig 5E). Moreover, while the levels of CCL11, CD40L, CXCL10, CXCL11, CXCL2, IFNα, IL6, and IL8 were significantly increased following bacterial stimulation in both lean and obese groups, the levels were higher in the obese group (Fig 5E).

Following viral ligand stimulation, fetal splenocytes from both lean and obese groups secreted CCL11, IFNγ, IL-8, IL-1β and PDGF-BB, with higher levels of the latter two analytes with pregravid obesity (Fig 5E). On the other hand, CCL20, CXCL11, CD40L, CXCL10, CXCL13, CXCL2, IFNα, IFNβ, IL-12, IL-13, IL-6, PDL1, TNFα and VEGF were exclusively induced by splenocytes from the obese group (Fig 5E). Following PMA stimulation, fetal splenocytes from both lean and obese groups generated a BDNF, CCL11, CCL2, CCL5, CD40L, CXCL2, GZMB, IL-8 and PDGF-AA response, with higher levels of the last two mediators in the obese group (Fig 5E). However, production of CCL20, CXCL13, GM-CSF, IFNβ, IL-13, PDL1,

PDGF-BB, and VEGF were significantly increased only in fetal splenocytes from the obese group, while CXCL11 and FGFβ were elevated exclusively in fetal splenocytes from lean controls (Fig 5E).

### Impact of Pregravid obesity on Transcriptional Changes in Fetal Splenocytes

We next assessed molecular mechanisms responsible for functional changes observed in splenic leukocytes using scRNAseq analysis. We identified 16 cell clusters within CD45+ splenic leukocytes, including pDC (*CD14-CD1c+SELL^hi^*), mDC (*CD14-CD1c+)*, macrophages (*MAMU-DRA+CD14^lo^LYZ+CD68+),* monocytes (*MAMU-DRA+CD14^hi^CD163+),* and HSPC (*CD34+)* (Fig 6A,B and Supp. Fig3E). The T cell compartment comprised of a proliferating cluster (*MKI67+*); two CD4 T cell clusters (*CD3+CD8-)* delineated based on of *CCR7* and *CD28* expression; a Treg (*IL2RA+CTLA4+)* cluster; and two CD8 T cell clusters (*CD3+CD8+*) separated based on granzymes (*GZMA, GZMB)* expression levels (Fig 6A,B). NK cells were defined as the *CD3-KLRB1+* population, and NKT cells were defined as the *CD3+ZBTB16+* population (Fig 6A,B). Three B cell clusters (*MS4A1+*) were identified, including Bcells_1 and Bcells_2 distinguished by differential expression of *CD1C* and *CD79A* (Fig. 6A,B). The third B cell population consisted of plasmablasts (SSR4^hi^XBP1^hi^) and was predominately enriched in the lean group (Fig. 6A-C and Supp. Fig3E,F). The loss of the plasmablast cluster was accompanied by a significant increase in the relative frequencies of Bcells_1 and Bcells_2 in the obese group (Fig 6C). Additionally, the frequencies of the CD4_1 and the CD8 T cell clusters were reduced with maternal pregravid obesity (Fig. 6C). Finally, pregravid obesity was associated with a modest increase in splenic macrophage population and a significant decrease in the frequency of HSPC (Fig. 6C).

**Figure 6:**
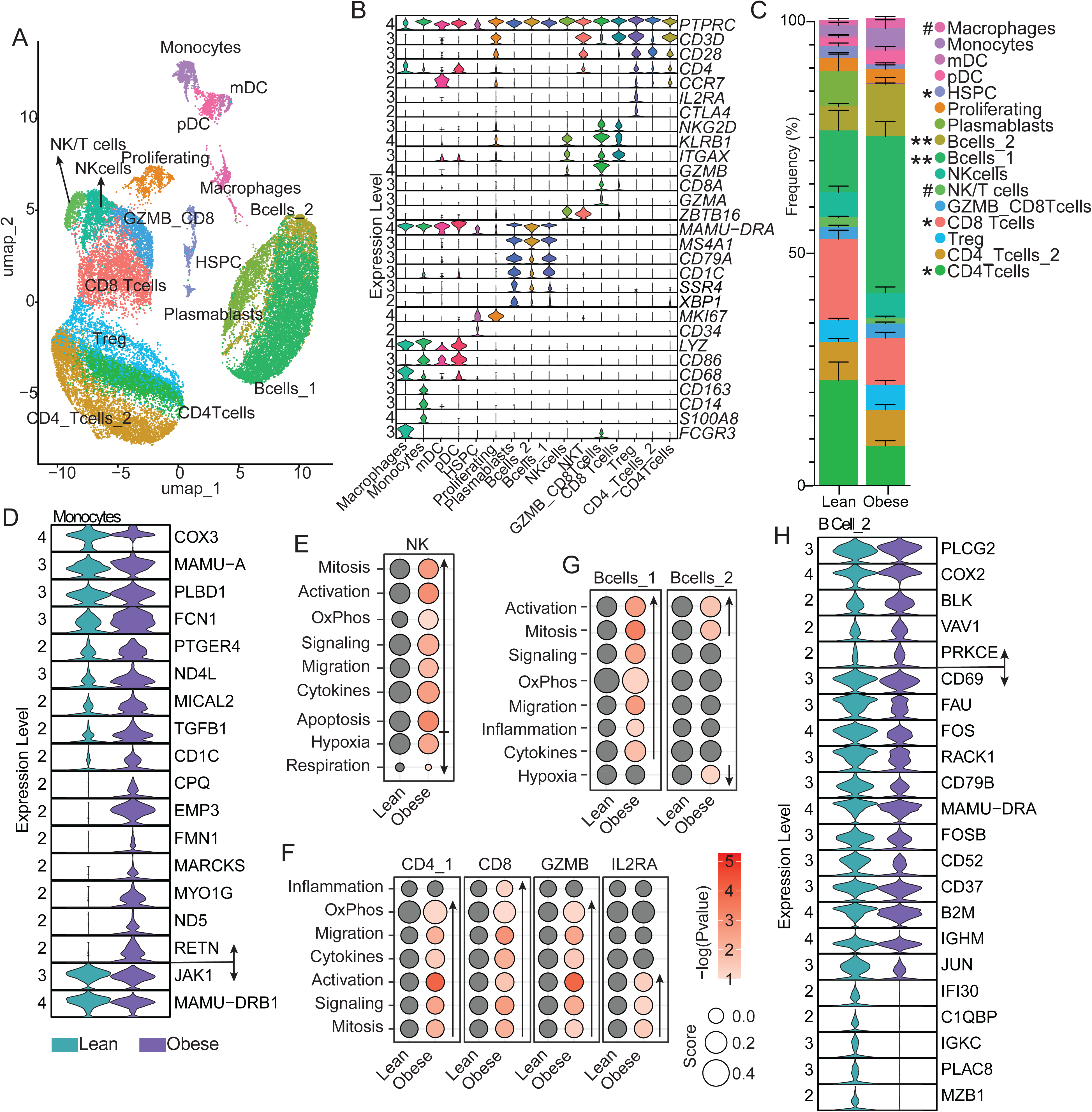
Impact of pregravid obesity on gene expression in fetal splenocytes. A) UMAP visualization of scRNAseq analysis of splenocytes, with colors indicating distinct cell clusters. B) Violin plots of marker genes used to define each cluster. C) Stacked bar plot showing relative cluster frequencies. #p < 0.1, *p < 0.05, **p < 0.01. D, H) Violin plots showing expression of the indicated differentially expressed genes (DEGs) within the indicated clusters. E–G) Bubble plots depicting module scores for the indicated clusters. Bubble size represents the average module score, while bubble color reflects the p-value from multiple t-test comparisons.

Although we did not detect any significant differences in the assessed module scores for splenic macrophages, we observed dysregulation of key genes in the obese group. Specifically, we noted a significant decrease in the expression of *JAK1*, a cytoplasmic tyrosine kinase essential for cellular responses and cytokine production, as well as a decrease in *MAMU-DRB1*, an MHC class II molecule required for antigen presentation (Fig 6D). Additionally, we observed upregulation of metabolic genes (*COX3, ND4L, PLBD1*), regulators of actin cytoskeleton involved in migration and phagocytosis (*EMP3, MICAL2, FMN1, MARCKS, MYO1G*), and pro-inflammatory genes (*FCN1, RETN*) (Fig 6D). Additionally, we observed increased expression of *TGFB1*, a profibrotic growth factor, and *CD1C*, a molecule involved in lipid antigen presentation to T cells in DC and whose expression in monocytes has been linked to a hyperinflammatory state (Fig 6D)(61, 62). In the macrophage subset, we observed increased scores associated with wound healing and OxPhos (Supp Fig 3G). We observed increased OxPhos concomitant with decreased hypoxia in mDCs, in addition to a decrease in cellular respiration in HSPC (Supp Fig 3G).

Fetal splenic lymphocyte subsets also exhibited changes following exposure to pregravid obesity. NK cells showed increased scores for mitosis, activation, OxPhos, cytokines, signaling, and migration modules (Fig 6E). Module scores for inflammation mitosis, activation, OxPhos, cytokines, signaling, and migration modules were increased to different extents across the T cells and B cell clusters (Fig 6F,G). Additionally, DEG analysis showed upregulation of genes essential for BCR signaling, including *PLCG2, BLK, PRKCE and VAV1* with pregravid obesity in B cells while key genes required for B cell maturation and germinal center formation, including *CD69, SOX4, CD79B, CD37, IGHM, and IGKC*, were downregulated (Fig 6H, Supp Table 2). We also observed downregulation of genes involved in antibody assembly, class switching, secretion, and plasma cell differentiation (*FAU, VAMP8, AIMP1, RHOB, and MZB1*) as well as inflammasome genes (*CASP1* and *AIM2*) and MHC class 1 molecule (*B2M*) (Fig 6H, Supp Table 2).

CellChat analysis identified a modest increase in the number and strength of potential signaling interactions in fetal splenocytes with pregravid obesity (Fig. 7A, B). We also identified several signaling pathways predicted to be stronger in the obese group. including CSF, RELN, EPHB, BAG, PDGF and IL2 whereas only CD39 signaling was elevated in the lean group (Fig. 7C). We next examined the differential use of signaling pathways across cell clusters between the two groups (Fig. 7D). Loss of CD39 signaling in the obese group was driven by reduced signaling from monocytes to multiple clusters (Fig. 7D,E). Signaling via TNF was increased from NKT and proliferating cells (Fig. 7D,F). Finally, PECAM1 and TGFb signaling from monocytes to multiple cell clusters was enhanced (Fig. 7D,G,H).

**Figure 7:**
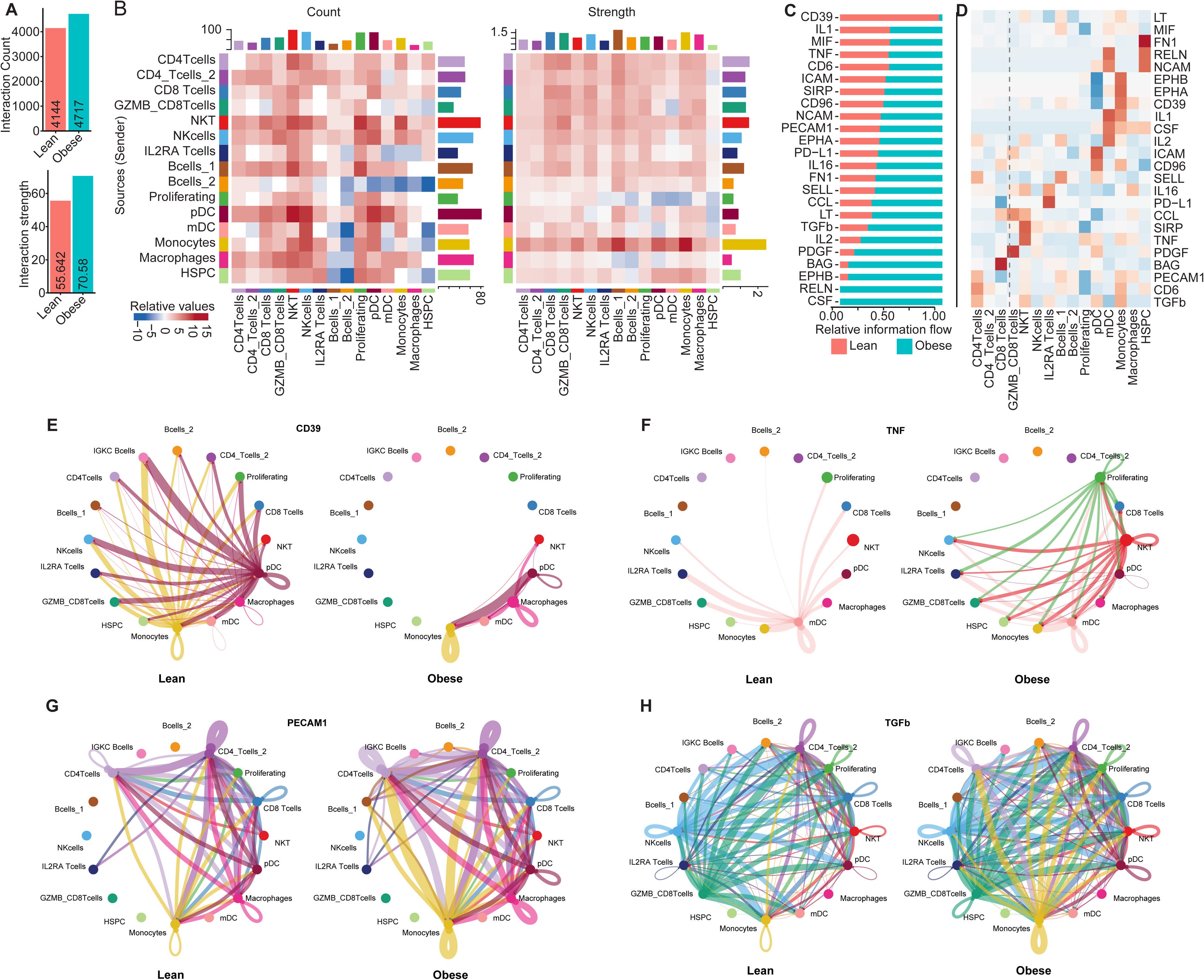
Pregravid obesity alters cell-cell communications in fetal splenocytes. A) Bar plot showing the inferred number and strength of cell–cell interactions. using CellChat analysis on the scRNAseq data. B) Heatmap depicting relative interaction counts and interaction strengths between clusters. Blue indicates reduced interactions in the obese group compared with the lean group. Bars along the top and right margins represent cumulative signaling interactions for the corresponding column and row, respectively. C) Bar plot showing relative information flow, defined as the aggregate probability of communication, for each signaling pathway. D) Heatmap illustrating differential cluster-to-pathway interactions. Red indicates increased interaction potential in the obese group relative to the lean group. E–H) Circle plots showing cluster-to-cluster interactions for the indicated signaling pathways. Lean samples are shown on the left and obese samples on the right. Line color denotes the source cell population, and line thickness reflects the relative interaction count.

### Immune Landscape and Functional Response of the Fetal Lung Leukocytes

Next, we assessed the immune landscape of the fetal lung. Spectral flow cytometry showed a significant increase in the frequencies of macrophages, mDC and pDC populations in fetal lung leukocytes exposed to pregravid obesity (Fig 8A,B, Supp Fig 4A). While the overall frequency of NK cells did not change, the frequency of the cytotoxic CD16+ NK cells was significantly increased (Fig 8C). We also observed a significant increase in the overall frequency of B cells with no change in the memory/naïve subset distribution (Fig 8D). Finally, we noted there was no change in the frequency of CD4 or CD8 T cells or their subsets. (Supp Fig 4B). Next, we examined the impact of pregravid obesity on the ability of fetal lung leukocytes to respond to stimulation. Interestingly, basal (NS) levels of multiple immune mediators, including CCL20, CD40L, CXCL11, CXCL13, IFNα, IL-12, IL-6, and VEGF were elevated with pregravid obesity (Fig 8F). Bacterial stimulation resulted in a more robust CCL11, CCL20, CD40L, CXCL11, CXCL13, CXCL2, and IL-12 response in the lean group (Fig 8F). In contrast, leukocytes exposed to pregravid obesity uniquely produced significant amounts of IFNγ, IL-10, IL-13, and IL-1β (Fig 8F). Only CCL2 was significantly produced following bacterial stimulation by both groups, and its production was further elevated in the pregravid obesity group (Fig 8F). Viral stimulation elicited a response driven by CXCL13 and IFNα in the lean group, while the response of the obese group was driven by IFNγ, IL-10, and IL-1β (Fig 8F). While leukocytes from both groups secreted GM-CSF, PD-L1, and TNFα in response to PMA stimulation, levels were higher with pregravid obesity (Fig 8F).

**Figure 8:**
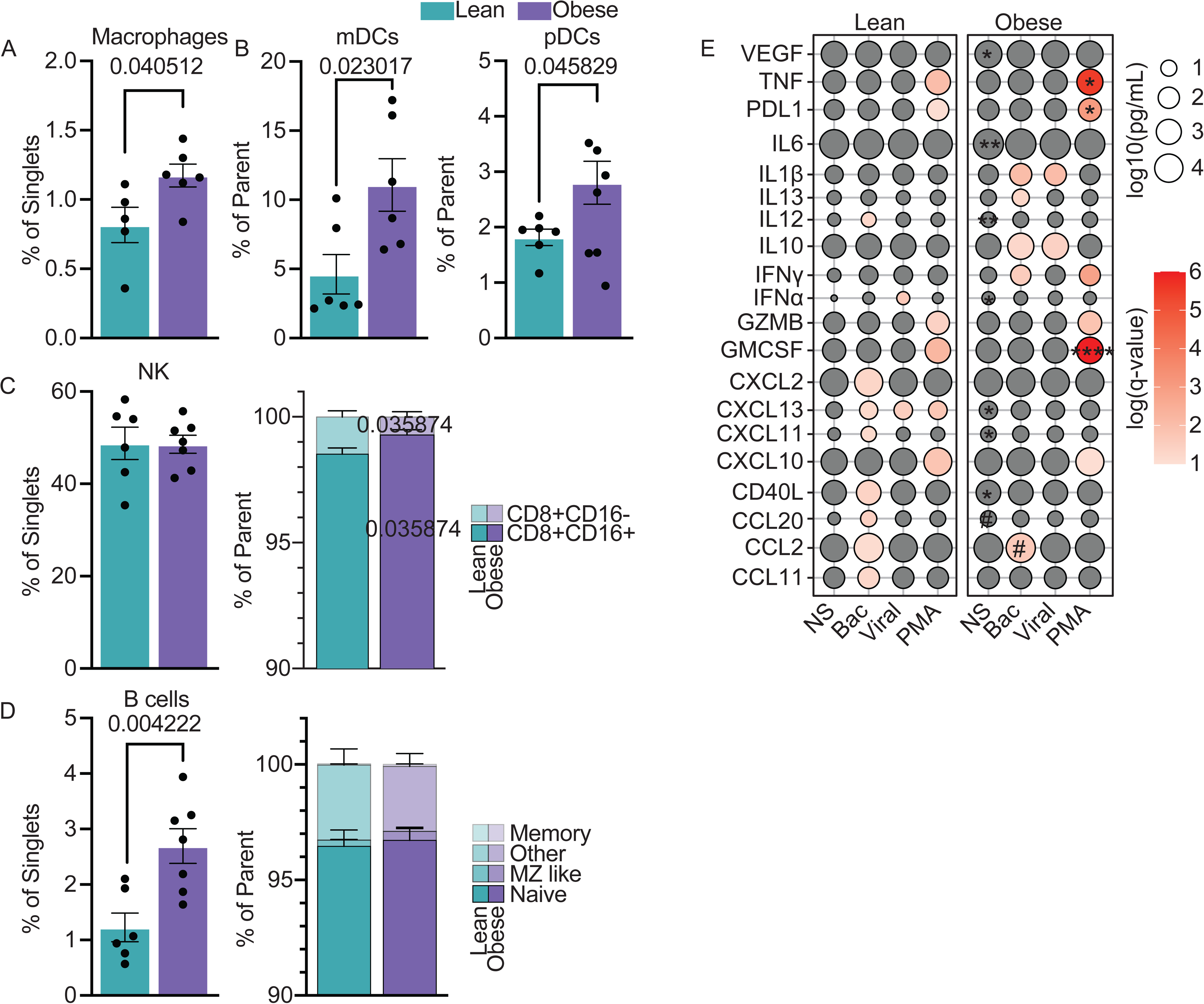
Pregravid obesity alters fetal lung immune cell composition. A–D) Bar plots showing the frequency of the indicated cell types in each group using spectral flow cytometry. The gating strategy is shown in Supplementary Fig. 4A. E) Bubble plot of cytokine production following ligand cocktail stimulation. Bubble size represents cytokine concentration log10(pg/mL), while bubble color reflects the p-value from a two-way ANOVA comparing stimulated conditions to the non-stimulated (NS) condition within each group. #p < 0.1, *p < 0.05, **p < 0.01 for comparisons of cytokine production after stimulation in obese relative to lean groups.

### Impact of Pregravid obesity on Transcriptional Responses in Fetal Lung Leukocytes

We used scRNA-seq to uncover mechanisms underlying altered functional responses by lung leukocytes. We identified alveolar macrophages (AM, *CD14^lo^MRC1+*), interstitial macrophages (IM, *CD14+MRC-ITGAM+),* and two subsets of infiltrating macrophages (Mac_1 and Mac_2) distinguished by differential expression of *CD68 and CD86* (Fig. 9A,B and Supp Fig 4C). The lymphoid compartment included two clusters of CD4 T cells (*CD3+KLRB1- and CD3+KLRB1-CD28+*), BCL11B high T cells (*CD3+KLRB1-BCL11B+),* CD8 T cells (*CD3+KLRB1+*), and a granzyme A high CD8 T cells (*CD3+KLRB1+GZMA+)* (Fig. 9A,B). NK cells were again identified as *CD3-KLRB1+,* whereas B cells as *MS4A1+* (Fig. 9A,B). Additionally, we identified a cluster of group 2 innate lymphoid cell (ILC2) characterized by the absence of *CD3* and positive for *IL7R, IL1RL1,* and *GATA3* (Fig. 9A,B). Pregravid obesity resulted in a significant decrease in the frequency of AM accompanied by a modest increase in Mac_1 (Fig. 9C). Additionally, we observed a significant decrease in CD4 T cells concomitant with a significant increase in NK and GZMA CD8 T cells (Fig. 9C).

**Figure 9:**
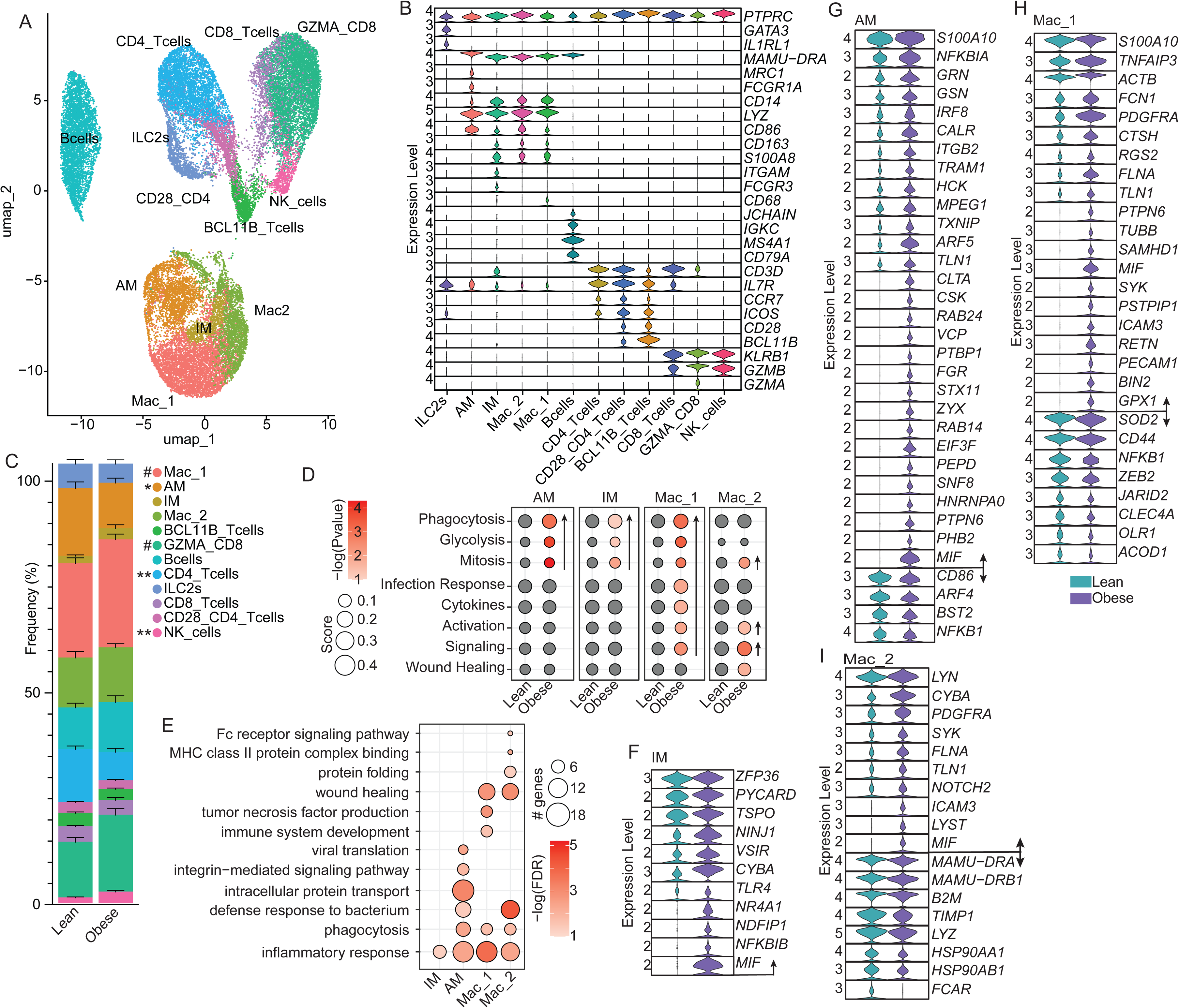
Impact of pregravid obesity on gene expression in fetal lung leukocytes. A) UMAP visualization of scRNA-seq lung leukocytes, with colors indicating distinct cell clusters. B) Violin plots of marker genes used to define each cluster. C) Stacked bar plot showing relative cluster frequencies. #p < 0.1, *p < 0.05, **p < 0.01. D) Bubble plot depicting module scores for the indicated clusters. Bubble size represents the average module score, while bubble color reflects the p-value from multiple t-test comparisons. E) Bubble plot of Gene Ontology (GO) terms enriched among differentially expressed genes (DEGs) in the indicated clusters. Bubble size represents the number of genes associated with each term, while bubble color indicates −log10(FDR). F–I) Violin plots showing expression of the indicated DEGs within the indicated clusters.

Changes in module scores and gene expression were restricted to innate cell clusters (Fig 9D). Module scores associated with phagocytosis and glycolysis were increased in AM, IM and the Mac_1 clusters with pregravid obesity (Fig. 9D). Both Mac_1 and Mac_2 showed increased activation and cell signaling scores (Fig. 9D). Mac_1 cluster exhibited an increase in scores related to cytokine production and the response to infections, while wound healing scores were higher in Mac_2 with pregravid obesity (Fig. 9D). IM cells exhibited modest transcriptional changes with pregravid obesity. DEGs in IM were primarily associated with inflammatory responses and cytokine production including *TLR4*, *CYBA*, *TSPO*, *MIF*, *NINJ1*, and *PYCARD* (Fig 9E,F). Additionally, the expression of multiple regulatory genes was also increased in IM with pregravid obesity, including *NFKBIB*, *NR4A1*, *NDFIP1*, and *SMURF1* (Fig 9E,F).

DEGs in AM cells mapped to processes related to inflammatory responses, phagocytosis, intracellular transport, and adhesion (Fig 9E). The expression of genes involved in cell migration (*ARF5, ITGB2, and TLN1*) and pathogen clearance (*CALR, CTLA, GSN, MPEG1, STX11, and NR4A1*) was increased (Fig. 9G). Many genes associated with macrophage activation and inflammation were also upregulated, including kinases critical for Fc receptor signaling (*CSK, FGR, and HCK*), transcriptions factor *IRF8*, cytokines (*MIF, S100A10*), and inflammasome adaptors (*PYCARD, TX*NIP) (Fig. 9G). Additionally, we observed increased expression of *VCP*, an ATPase that regulates protein degradation and autophagy, and *VSIR*, an immune checkpoint molecule that inhibits macrophage-mediated T cell activation.

DEG in the Mac_1 cluster were associated with immune system development, cell killing, wound healing, and inflammatory responses (Fig 9E). We observed the downregulation of genes linked to antimicrobial defenses (*ACOD1*), macrophage polarization (*JARID2*), mitochondrial antioxidant activity (*SOD2*), and macrophage tissue-resident programming (*ZEB2*). Upregulated genes were involved in phagocytosis(*CTSH, NR4A1*), pattern recognition (*FCN1, PTPN6*), adhesion and migration (*ICAM3, PECAM1, FLNA, PSTPIP1, RGS2* ), oxidative stress (*GPX1, SYK*), and proinflammatory cytokine production (*MIF, S100A10, RETN*) (Fig. 9H). DEGs in the Mac_2 cluster mapped to inflammatory responses, cell killing and wound healing as well as protein folding, MHC class II function, degranulation, and Fc receptor signaling (Fig 9E). Genes essential for antigen processing and presentation to T cells (*MAMU-DRA, MAMU-DRB, B2M*), immune receptor assembly (*HSP90AA1, HSP90AB1, HYPK, PPIB*), bactericidal activity (*LYZ, FCAR*), and anti-inflammatory signaling (*ADORA2B*) were downregulated. (Fig 9I). On the other hand, genes important for migration/adhesion (*FLNA, ICAM3, PECAM1, TLN1*), phagocytic receptor signaling (*SYK, LYN, LYST*), inflammation (*MIF*, *PDGFRA*) were upregulated (Fig 9I).

As described for the spleen, the number and strength of potential interactions in fetal lung leukocytes was increased with pregravid obesity (Fig. 10A). Signaling alog TGFb, IL16, CD40, ICAM, MIF, and PROS pathways was predominantly upregulated in the innate cell clusters, including with pregravid obesity (Fig. 10B,C). In contrast, CD39 and CCL signaling was upregulated exclusively in the lean group (Fig. 10C). Interestingly, the IM cluster exhibited the largest changes in signaling pathways with increased signaling via VEGF, SELPLG, IL16, FN1, and TGFb (Fig. 10D,E). As described for UCBMC and spleen, CD39 signaling was severely decreased with exposure to pregravid obesity, especially in the AM and IM clusters (Fig. 10D,F). With pregravid obesity, ICAM saw a loss of signaling in the AM cluster while CD4 T cells emerged as a new source of signaling to multiple clusters, and ILC2s were recipient of ICAM signaling with pregravid obesity (Fig. 10D,G). Finally, GZMA_CD8 T cells and B cells became new sources of MIF signaling with pregravid obesity (Fig. 10D,H).

**Figure 10:**
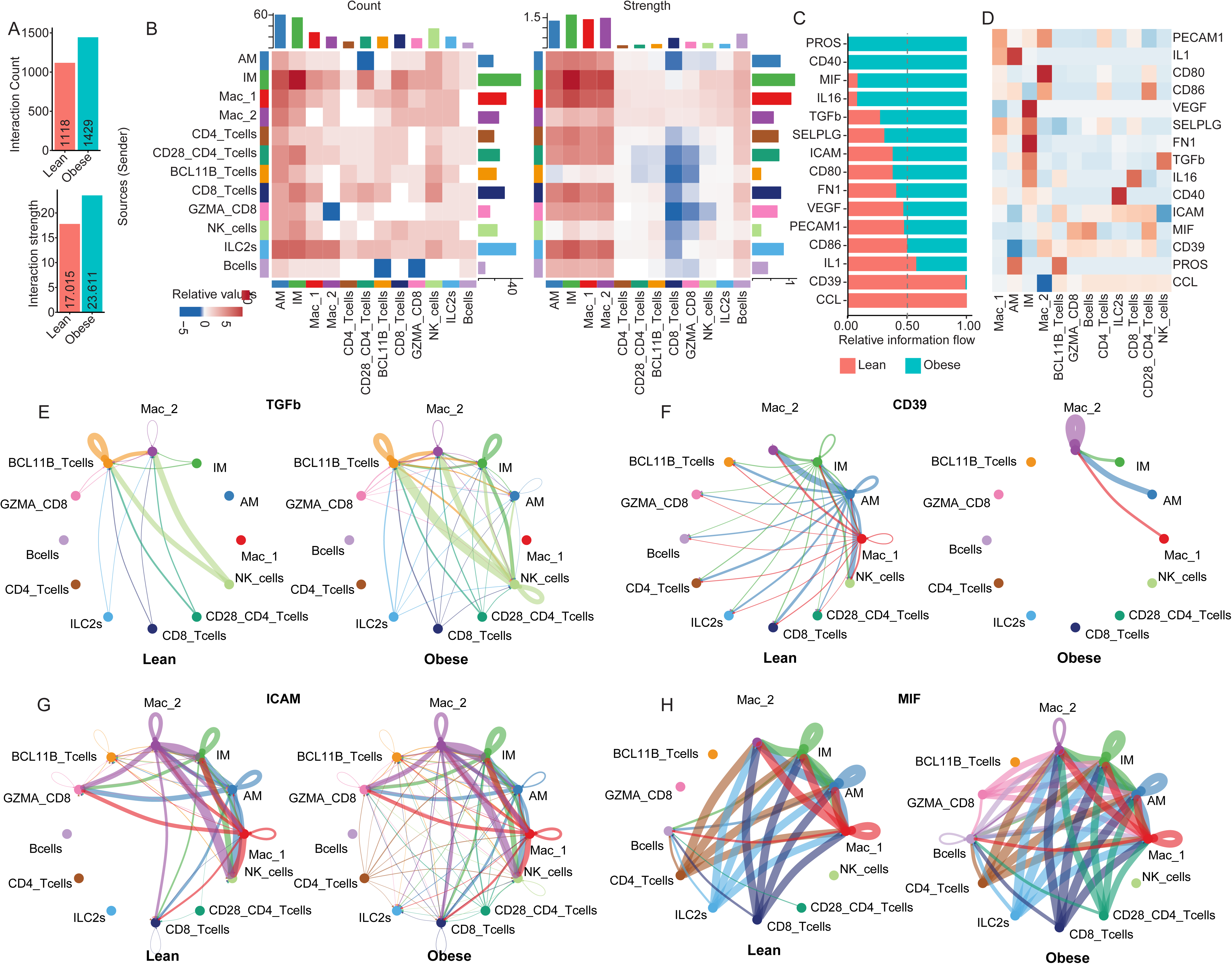
Pregravid obesity alters cell-cell communications in fetal lung leukocytes. A) Bar plot showing inferred interactions and their strength using CellChat analysis of scRNA-seq data. B) Heatmap depicting the relative count and interaction strength between clusters. Blue indicates reduced interaction in the obese group compared to the lean group. Bars along the top or right side represent the cumulative interaction of all signaling for the corresponding column or row, respectively. C) Bar plot of relative information flow, representing the aggregate probability of communication for each signaling pathway. D) Heatmap showing differential relative cluster-to-pathway interactions. Red indicates higher interaction potential in the obese group compared to the lean group. E–H) Circle plots illustrating cluster-to-cluster interactions for the specified pathways. The lean group is shown on the left and the obese group on the right. Line color represents the source of the interaction, and line thickness corresponds to the relative interaction count.

## Discussion

Pregravid obesity has been linked to adverse health outcomes for offspring, including increased susceptibility to infections, asthma, and metabolic disorders(7-19). Recent findings indicate that pregravid obesity alters immune cell development, frequency and functional capacity (20-24, 27, 28). Clinical studies are inherently limited in their ability to assess mechanistic effects due to restricted access to fetal tissues. Existing animal models have overly relied on rodent models and obesogenic diets, which may not fully recapitulate human obesity(36-39). To address this knowledge gap, we examined the impact of spontaneous pregravid obesity on circulating and tissue-resident leukocytes in fetal UCB, spleen, and lung, using a nonhuman primate model of spontaneous obesity. Our findings reveal that pregravid obesity fundamentally reshapes the developing fetal immune landscape across both lymphoid and non-lymphoid tissues.

The NHP model of spontaneous obesity recapitulates many metabolic and inflammatory features characteristic of human pregravid obesity and aligns with previous observations in NHP models of diet-induced obesity including impaired glucose tolerance, hyperinsulinemia, and elevated fasting glucose (46, 47). The increased placental weight observed in the obese group aligns with previous studies showing altered placental morphology, increased placental lipid accumulation, heightened placental inflammation, and disrupted nutrient transport with pregravid obesity(63-65). The increase in fetal heart and lung weight, coupled with decreased pancreatic and adrenal gland weight, indicates organ-specific developmental reprogramming. Indeed, excess inflammation, which we observed transcriptionally and functionally, has been associated with reduced lung compliance and capacity, and impaired gas exchange(66, 67). Additionally, pregravid obesity has been associated with increased rates of asthma and severe RSV infection(17, 19). The increase in heart weight has been previously observed in mouse models of pregravid obesity(68). Furthermore, we observed elevated expression and signaling of *TGFB1* and *RETN* genes in multiple clusters within splenocytes and lung leukocytes in the obese group. TGFβ promotes collagen accumulation in fetal myocardium, inducing fibrosis and leading to increased wall thickness and hypertrophy(69), while RETN (Resistin) promotes cardiac hypertrophy by increasing cardiomyocyte cell size(70). In contrast, fetal pancreas was decreased in size. This finding aligns with previous rodent studies showing that pregravid obesity leads to decreased beta-cell mass with altered pancreatic morphology and structure throughout early and adult life(71, 72). The pattern of organ-specific growth suggests that tissues with substantial mesodermal contributions may be particularly sensitive to pregravid obesity. Interestingly, the immune system is predominantly mesoderm-derived(73).

Functional assessment revealed hyperresponsiveness to bacterial, viral, and PMA stimulation in all three compartments. Splenic leukocytes showed increased production of type I interferons (IFNα, IFNβ) and chemokines (CXCL10, CXCL11). However, the simultaneous increased production of regulatory cytokines (IL-10) indicates attempts to dampen excessive inflammation. Lung leukocytes showed excess basal cytokine production. Increased levels of VEGF production have been associated with airway wall thickening, airway edema and obstruction(74) in line with increased respiratory disease in offspring of women with obesity (17, 19). Additionally, levels of IL-1β in sputum, bronchoalveolar lavage fluid, and serum of asthma patients correlates with disease severity and poor lung function(75). The observed increase in response to stimulation contrasts with our previous reports of blunted monocyte responses in term human UCB(23) and dampened responses to E. Coli stimulation by splenic and ileal macrophages in an NHP model of WSD-induced obesity(23), but are aligned with data from rodent models of maternal HFD(25, 26).

We recently showed, using the same cohort of fetuses, that pregravid obesity is associated with a transcriptional shift towards myelopoiesis in HSPCs (29). The monocyte progenitors in the fetal bone marrow (FBM) also showed markers consistent with possible impairment of monocyte egress from the bone marrow (29). In line with these findings, we observed a decrease in the total monocyte frequency in UCB from spontaneously obese dams.

This decreased total monocyte frequency was also observed in human UCB with pregravid obesity(23). In contrast, the frequency of macrophages in spleen and lung tissue was significantly increased with pregravid obesity. Tissue-resident macrophages are seeded throughout gestation with primitive yolk sac-derived erythro-myeloid progenitors, followed by fetal liver monocytes during mid-gestation, and finally by bone marrow-derived monocytes during late gestation and postnatally(76-78). The decreased frequency of tissue resident alveolar macrophages coupled with the increase in monocyte-derived infiltrating macrophages has been shown to transform the lung from an immunologically quiescent organ to an inflammatory one, which aligns with increased offspring respiratory disease with pregravid obesity(79). We also observed a shift from MHC class II (*MAMU-DRB1*) to MHC class I (*MAMU-A*) expression in multiple tissues with pregravid obesity. Downregulation of MHC class II expression in myeloid cells lead to defective antigen presentation, resulting in impaired CD4+ T cell activation and a shift toward dysfunctional states(80).

Fetal lung macrophages exhibited the most complex transcription alterations. Within the Mac_1 populations, the downregulation of ACOD1, an inflammation-induced enzyme that generates itaconate and regulates macrophage metabolic and inflammatory responses(81), may contribute to the hyperresponsiveness we observed in the present study. The downregulation of *JARID2*, an epigenetic regulator that controls macrophage polarization, and *SOD2*, a mitochondrial antioxidant enzyme that detoxifies superoxide radicals, suggests compromised ability to resolve inflammation(82). Additionally, adenosine generated by CD39 signaling is critical for maintaining tolerance to inhaled antigens and limiting excessive inflammation during infections in the lung (83, 84). Therefore, its downregulation is in line with increased incidence of asthma and wheezing detected in offspring of women with obesity.

In the spleen, We observed increased TGFβ production, which has been shown to promote fibrosis and contribute to sustained tissue inflammation(85). Additionally, we observed increased CD1C expression in splenic macrophages. CD1C presents lipid antigens to T cells and has been associated with hyperinflammatory states as well as the differentiation to more hyperresponsive DC subsets(86). CD1C upregulation may reflect altered lipid metabolism in the fetal spleen. In splenic macrophages with pregravid obesity, we also observed decreased *JAK1* expression, which is essential for anti-inflammatory cytokine signaling, therefore impairing the ability to regulate the inflammatory state(87).

The upregulation of OxPhos genes in UCBMC alongside activation markers suggests metabolic stress in T cells(88). However, we also observed the downregulation of genes involved in T cell function and identity. The downregulation of critical TCR components (*CD3G*) and lineage-defining transcription factors (*RUNX1, BCL11B)* could compromise T cell activation (89, 90). Indeed, previous studies have reported poor responses to stimulation by T cells and poor vaccine responses in early life with pregravid obesity (23, 30-35). Additionally, in UCBMC CD8 T cells, we saw upregulation of components of the mitochondrial electron transport chain (*COX3, COX5A*) and *TGFB1*, which together with downregulation of several genes encoding cytotoxic molecules suggest a compromised function. Additionally, we observed the premature shift of spleen CD4 T cells toward terminal effector memory (TEM), potentially explaining epidemiological observations of poor vaccine responses and increased infection susceptibility in offspring of mothers with obesity(17, 19).

The frequencies of B cells were significantly increased in all fetal tissues with pregravid obesity. We recently reported a significantly decreased frequency of lymphoid progenitors in the bone marrow(29), suggesting acceleration of B cell progenitor trafficking to the periphery. B cells exhibited decreased isotype class switching module scores together with reduced expression of *IGKC*. Given that 60-70% of B cells express kappa light chains, this finding suggests compromised antibody production capacity(91). Upregulation of *FCRL2* and *PTPRJ*, both of which attenuate BCR signaling, indicates premature differentiation or functional exhaustion(92). We also observed dysregulation of several genes that regulate chemokine signaling and cell migration suggesting that pregravid obesity alters B cells trafficking.

In the spleen, we observed the complete loss of the plasmablast population. Plasmablasts and plasma cells produce the IgM antibodies that constitute neonate’s first line of endogenous humoral defense prior to the maturation of germinal center-dependent responses, with the fetal splenic compartment serving as a key extrafollicular niche for their early differentiation from B1 cell precursors derived from the fetal liver(93-95). In adults with obesity, preferential B cell activation through the extrafollicular pathway has been shown to deplete germinal center precursors and reduces the generation of plasma cells(96). The downregulation of plasma cell differentiation markers and components of antibody assembly machinery indicate a block in plasma cell differentiation. The downregulation of inflammasome components further compromises B cell function. Loss of CD40 signaling, critical for T cell-B cell interactions and germinal center formation, may further exacerbate B cell dysfunction. The inability to generate appropriate antibody responses, particularly class-switched high-affinity antibodies, would leave offspring vulnerable to bacterial and viral pathogens in early life.

This study demonstrates that spontaneous pregravid pregravid obesity, independent of dietary intervention, fundamentally reprograms the developing fetal immune system in a coordinated manner across multiple tissues. We identified a consistent pattern of innate immune priming with hyperresponsiveness to stimulation, impaired maturation in B cell and T cell compartments, altered tissue distributions, and profound transcriptional reprogramming affecting antigen presentation, cytokine signaling, and loss of tolerogenic signals. These alterations create a pro-inflammatory environment that contribute to increased incidence of sepsis, severe respiratory infections, poor vaccine responses and increased autoimmune disease risk in offspring of mothers with obesity(7-19). Understanding these mechanisms is critical for developing interventions to protect offspring health and break the intergenerational cycle of obesity-associated disease. Future studies should examine earlier gestational timepoints and expand to interrogate the fetal yolk sac and liver to determine when immune alterations first emerge and whether they coincide with specific developmental windows.

## Acknowledgments

We thank members of the veterinary, pathology and surgical staff at the ONPRC for support with animal studies. This work was supported by the UK Flow Cytometry & Immune Monitoring core facility. (RRIDSCR_026358).

## Author contributions

O.V. and I.M. conceived the idea. L.D.M and T.H. worked with the animal model. O.V. and U.A. collected tissue samples. B.M.D., S.B.W., O.V. and U.A. processed tissue samples. B.M.D. and S.B.W completed all experiments. B.M.D. performed the analysis. B.M.D., O.V., and I.M. interpreted the data and prepared the manuscript. All authors read and approved the final manuscript.

## Data Availability

Data underlying the transcriptome analysis is available in the SRA under project number PRJNA1247568.

---

**Supp Figure 1: Metabolic parameters and maternal immune mediator levels.**

A-B) Line graph of dams’ fasting glucose and insulin levels throughout gestation (left) and bar graphs of corresponding AUC values (right).

C) Bubble plot of AUC analysis for cytokine levels in maternal plasma. Bubble size represents average AUC, while bubble color reflects the p-value from the multiple t-tests comparing lean to obese samples.

**Supp Figure 2: UCBMC composition and analysis.**

A) Gating strategy of UCBMC spectral flow cytometry.

B-C) bar plots of indicated cell frequencies in UCBMC for lean (left) and obese (right) groups.

D) UMAP colored by group origin.

E-G) Modules scores for the indicated cell clusters from UCBMC for lean (left) and obese (right) groups.

**Supp Figure 3: Fetal spleen immune cell composition and analysis.**

A) Gating strategy of spleen leukocyte spectral flow cytometry.

B-D) bar plots of indicated cell frequency.

E) UMAP colored by group origin.

F) Bubbleplot of GO terms for marker genes of Bcell clusters. Size of the bubble indicates the number of genes mapping to the term and color of the bubble indicates -log10(qvalue).

G) Modules scores for the indicated cluster.

**Supp Figure 4: Fetal lungs immune cell composition and analysis.**

A) Gating strategy of fetal lung spectral flow cytometry.

B) bar plots of indicated cell frequency.

C) UMAP colored by group origin.

Supp Table 1: Marker genes for scRNAseq clusters

Supp Table 2: DEGs for scRNAseq clusters

Supp Table 3: Organ weights.

## References

1. 1. Emmerich SD FC, Stierman B, Ogden CL. Obesity and severe obesity prevalence in adults: United States, August 2021–August 2023. National Center for Health Statistics. 2024.

2. 2. Wang Y, Beydoun MA, Min J, Xue H, Kaminsky LA, Cheskin LJ. Has the prevalence of overweight, obesity and central obesity levelled off in the United States? Trends, patterns, disparities, and future projections for the obesity epidemic. International Journal of Epidemiology. 2020;49(3):810–23.

2. Driscoll AK, Gregory ECW. Increases in Prepregnancy Obesity: United States, 2016-2019. NCHS Data Brief. 2020(392):1–8.

3. Leung TY, Leung TN, Sahota DS, Chan OK, Chan LW, Fung TY, et al. Trends in maternal obesity and associated risks of adverse pregnancy outcomes in a population of Chinese women. BJOG. 2008;115(12):1529–37.

4. Haque R, Keramat SA, Rahman SM, Mustafa MUR, Alam K. Association of maternal obesity with fetal and neonatal death: Evidence from South and South-East Asian countries. PLoS One. 2021;16(9):e0256725.

5. van Duijn L, Rousian M, Hoek J, Willemsen SP, van Marion ES, Laven JSE, et al. Higher preconceptional maternal body mass index is associated with faster early preimplantation embryonic development: the Rotterdam periconception cohort. Reprod Biol Endocrinol. 2021;19(1):145.

6. Catalano PM. Increasing maternal obesity and weight gain during pregnancy: the obstetric problems of plentitude. Obstet Gynecol. 2007;110(4):743–4.

7. Gaillard R, Felix JF, Duijts L, Jaddoe VW. Childhood consequences of maternal obesity and excessive weight gain during pregnancy. Acta Obstet Gynecol Scand. 2014;93(11):1085–9.

8. Gupta A, Singh A, Fernando RL, Dharmage SC, Lodge CJ, Waidyatillake NT. The association between sugar intake during pregnancy and allergies in offspring: a systematic review and a meta-analysis of cohort studies. Nutr Rev. 2021.

9. Wilson RM, Marshall NE, Jeske DR, Purnell JQ, Thornburg K, Messaoudi I. Maternal obesity alters immune cell frequencies and responses in umbilical cord blood samples. Pediatr Allergy Immunol. 2015;26(4):344–51.

10. Ardura-Garcia C, Cuevas-Ocana S, Freitag N, Kampouras A, King JA, Kouis P, et al. ERS International Congress 2020: highlights from the Paediatric Assembly. ERJ Open Res. 2021;7(1).

11. Godfrey KM, Reynolds RM, Prescott SL, Nyirenda M, Jaddoe VW, Eriksson JG, et al. Influence of maternal obesity on the long-term health of offspring. Lancet Diabetes Endocrinol. 2017;5(1):53–64.

12. Sharp GC, Salas LA, Monnereau C, Allard C, Yousefi P, Everson TM, et al. Maternal BMI at the start of pregnancy and offspring epigenome-wide DNA methylation: findings from the pregnancy and childhood epigenetics (PACE) consortium. Hum Mol Genet. 2017;26(20):4067–85.

13. Briese V, Voigt M, Hermanussen M, Wittwer-Backofen U. Morbid obesity: pregnancy risks, birth risks and status of the newborn. Homo. 2010;61(1):64–72.

14. Castro LC, Avina RL. Maternal obesity and pregnancy outcomes. Curr Opin Obstet Gynecol. 2002;14(6):601–6.

15. Danieli-Gruber S, Maayan-Metzger A, Schushan-Eisen I, Strauss T, Leibovitch L. Outcome of preterm infants born to overweight and obese mothers. J Matern Fetal Neonatal Med. 2017;30(4):402–5.

16. Griffiths PS, Walton C, Samsell L, Perez MK, Piedimonte G. Maternal high-fat hypercaloric diet during pregnancy results in persistent metabolic and respiratory abnormalities in offspring. Pediatr Res. 2016;79(2):278–86.

17. Rastogi S, Rojas M, Rastogi D, Haberman S. Neonatal morbidities among full-term infants born to obese mothers. J Matern Fetal Neonatal Med. 2015;28(7):829–35.

18. Suk D, Kwak T, Khawar N, Vanhorn S, Salafia CM, Gudavalli MB, et al. Increasing maternal body mass index during pregnancy increases neonatal intensive care unit admission in near and full-term infants. J Matern Fetal Neonatal Med. 2016;29(20):3249–53.

19. Friedman JE. Developmental Programming of Obesity and Diabetes in Mouse, Monkey, and Man in 2018: Where Are We Headed? Diabetes. 2018;67(11):2137–51.

20. Sureshchandra S, Mendoza N, Jankeel A, Wilson RM, Marshall NE, Messaoudi I. Phenotypic and Epigenetic Adaptations of Cord Blood CD4+ T Cells to Maternal Obesity. Front Immunol. 2021;12:617592.

21. Sureshchandra S, Wilson RM, Rais M, Marshall NE, Purnell JQ, Thornburg KL, et al. Maternal Pregravid Obesity Remodels the DNA Methylation Landscape of Cord Blood Monocytes Disrupting Their Inflammatory Program. J Immunol. 2017;199(8):2729–44.

22. Sureshchandra S, Doratt BM, Mendza N, Varlamov O, Rincon M, Marshall NE, et al. Maternal obesity blunts antimicrobial responses in fetal monocytes. Elife. 2023;12.

23. Gonzalez-Espinosa LO, Montiel-Cervantes LA, Guerra-Marquez A, Penaflor-Juarez K, Reyes-Maldonado E, Vela-Ojeda J. Maternal obesity associated with increase in natural killer T cells and CD8+ regulatory T cells in cord blood units. Transfusion. 2016;56(5):1075–81.

24. Odaka Y, Nakano M, Tanaka T, Kaburagi T, Yoshino H, Sato-Mito N, et al. The influence of a high-fat dietary environment in the fetal period on postnatal metabolic and immune function. Obesity (Silver Spring). 2010;18(9):1688–94.

25. Myles IA, Fontecilla NM, Janelsins BM, Vithayathil PJ, Segre JA, Datta SK. Parental dietary fat intake alters offspring microbiome and immunity. J Immunol. 2013;191(6):3200–9.

26. Sureshchandra S, Chan CN, Robino JJ, Parmelee LK, Nash MJ, Wesolowski SR, et al. Maternal Western-style diet remodels the transcriptional landscape of fetal hematopoietic stem and progenitor cells in rhesus macaques. Stem Cell Reports. 2022;17(12):2595–609.

27. Nash MJ, Dobrinskikh E, Soderborg TK, Janssen RC, Takahashi DL, Dean TA, et al. Maternal diet alters long-term innate immune cell memory in fetal and juvenile hematopoietic stem and progenitor cells in nonhuman primate offspring. Cell Rep. 2023;42(4):112393.

28. Doratt BM, Hemati H, Wagner SB, Blanton MB, Avila U, Varlamov O, et al. Maternal Obesity Reprograms Differentiation Trajectories of Fetal Hematopoietic Stem and Progenitor Cells Through Altered Inflammatory Signaling. bioRxiv. 2026:2026.01.29.702548.

29. Grayson BE, Levasseur PR, Williams SM, Smith MS, Marks DL, Grove KL. Changes in melanocortin expression and inflammatory pathways in fetal offspring of nonhuman primates fed a high-fat diet. Endocrinology. 2010;151(4):1622–32.

30. Li S, Kievit P, Robertson AK, Kolumam G, Li X, von Wachenfeldt K, et al. Targeting oxidized LDL improves insulin sensitivity and immune cell function in obese Rhesus macaques. Mol Metab. 2013;2(3):256–69.

31. McCurdy CE, Bishop JM, Williams SM, Grayson BE, Smith MS, Friedman JE, et al. Maternal high-fat diet triggers lipotoxicity in the fetal livers of nonhuman primates. J Clin Invest. 2009;119(2):323–35.

32. Wesolowski SR, Mulligan CM, Janssen RC, Baker PR, 2nd, Bergman BC, D’Alessandro A, et al. Switching obese mothers to a healthy diet improves fetal hypoxemia, hepatic metabolites, and lipotoxicity in non-human primates. Mol Metab. 2018;18:25–41.

33. Aagaard-Tillery KM, Grove K, Bishop J, Ke X, Fu Q, McKnight R, et al. Developmental origins of disease and determinants of chromatin structure: maternal diet modifies the primate fetal epigenome. J Mol Endocrinol. 2008;41(2):91–102.

34. Farley D, Tejero ME, Comuzzie AG, Higgins PB, Cox L, Werner SL, et al. Feto-placental adaptations to maternal obesity in the baboon. Placenta. 2009;30(9):752–60.

35. Grzeda E, Matuszewska J, Ziarniak K, Gertig-Kolasa A, Krzysko-Pieczka I, Skowronska B, et al. Animal Foetal Models of Obesity and Diabetes - From Laboratory to Clinical Settings. Front Endocrinol (Lausanne). 2022;13:785674.

36. Reynolds CM, Segovia SA, Vickers MH. Experimental Models of Maternal Obesity and Neuroendocrine Programming of Metabolic Disorders in Offspring. Front Endocrinol (Lausanne). 2017;8:245.

37. Rajamoorthi A, LeDuc CA, Thaker VV. The metabolic conditioning of obesity: A review of the pathogenesis of obesity and the epigenetic pathways that "program" obesity from conception. Front Endocrinol (Lausanne). 2022;13:1032491.

38. D’Angelo CS, Koiffmann CP. Copy number variants in obesity-related syndromes: review and perspectives on novel molecular approaches. J Obes. 2012;2012:845480.

39. Coe CL, Lubach GR. Maternal determinants of gestation length in the rhesus monkey. Trends Dev Biol. 2021;14:63–72.

40. Roberts VHJ, Castro JN, Wessel BM, Conrad DF, Lewis AD, Lo JO. Rhesus macaque fetal and placental growth demographics: A resource for laboratory animal researchers. Am J Primatol. 2023;85(8):e23526.

41. Summers L, Clingerman KJ, Yang X. Validation of a body condition scoring system in rhesus macaques (Macaca mulatta): assessment of body composition by using dual-energy X-ray absorptiometry. J Am Assoc Lab Anim Sci. 2012;51(1):88–93.

42. Clark AT, Gkountela S, Chen D, Liu W, Sosa E, Sukhwani M, et al. Primate Primordial Germ Cells Acquire Transplantation Potential by Carnegie Stage 23. Stem Cell Reports. 2017;9(1):329–41.

43. Burwitz BJ, Yusova S, Robino JJ, Takahashi D, Luo A, Slayden OD, et al. Western-style diet in the presence of elevated circulating testosterone induces adipocyte hypertrophy without proinflammatory responses in rhesus macaques. Am J Reprod Immunol. 2023;90(4):e13773.

44. Messaoudi I, Handu M, Rais M, Sureshchandra S, Park BS, Fei SS, et al. Long-lasting effect of obesity on skeletal muscle transcriptome. BMC Genomics. 2017;18(1):411.

45. Bishop CV, Stouffer RL, Takahashi DL, Mishler EC, Wilcox MC, Slayden OD, et al. Chronic hyperandrogenemia and western-style diet beginning at puberty reduces fertility and increases metabolic dysfunction during pregnancy in young adult, female macaques. Hum Reprod. 2018;33(4):694–705.

46. Bishop CV, Takahashi D, Mishler E, Slayden OD, Roberts CT, Hennebold J, et al. Individual and combined effects of 5-year exposure to hyperandrogenemia and Western-style diet on metabolism and reproduction in female rhesus macaques. Hum Reprod. 2021;36(2):444–54.

47. Jin S, Guerrero-Juarez CF, Zhang L, Chang I, Ramos R, Kuan CH, et al. Inference and analysis of cell-cell communication using CellChat. Nat Commun. 2021;12(1):1088.

48. Elhddad AS, Fairlie F, Lashen H. Impact of gestational weight gain on fetal growth in obese normoglycemic mothers: a comparative study. Acta Obstet Gynecol Scand. 2014;93(8):771–7.

49. Straughen JK, Trudeau S, Misra VK. Changes in adipose tissue distribution during pregnancy in overweight and obese compared with normal weight women. Nutr Diabetes. 2013;3(8):e84.

50. Hansen BC. The evolution of diabetes in non human primates: comparative physiology implications for human type 2 diabetes mellitus (T2DM). The FASEB Journal. 2010;24(S1):1055.10-.10.

51. Li MO, Wan YY, Sanjabi S, Robertson AK, Flavell RA. Transforming growth factor-beta regulation of immune responses. Annu Rev Immunol. 2006;24:99–146.

52. Thomas DA, Massague J. TGF-beta directly targets cytotoxic T cell functions during tumor evasion of immune surveillance. Cancer Cell. 2005;8(5):369–80.

53. Prazma CM, Yazawa N, Fujimoto Y, Fujimoto M, Tedder TF. CD83 expression is a sensitive marker of activation required for B cell and CD4+ T cell longevity in vivo. J Immunol. 2007;179(7):4550–62.

54. Glaser J, Neumann MH, Mei Q, Betz B, Seier N, Beyer I, et al. Macrophage capping protein CapG is a putative oncogene involved in migration and invasiveness in ovarian carcinoma. Biomed Res Int. 2014;2014:379847.

55. Li S, Chen A, Gui J, Zhou H, Zhu L, Mi Y. TLN1: an oncogene associated with tumorigenesis and progression. Discov Oncol. 2024;15(1):716.

56. Moratz C, Hayman JR, Gu H, Kehrl JH. Abnormal B-cell responses to chemokines, disturbed plasma cell localization, and distorted immune tissue architecture in Rgs1-/- mice. Mol Cell Biol. 2004;24(13):5767–75.

57. Shea LK, Honjo K, Redden DT, Tabengwa E, Li R, Li FJ, et al. Fc receptor-like 2 (FCRL2) is a novel marker of low-risk CLL and refines prognostication based on IGHV mutation status. Blood Cancer J. 2019;9(6):47.

58. Delespesse G, Hofstetter H, Sarfati M. Low-affinity receptor for IgE (FcERII, CD23) and its soluble fragments. Int Arch Allergy Appl Immunol. 1989;90 Suppl 1:41–4.

59. Pinsky DJ. Cd39 as a Critical Ectonucleotidase Defense against Pathological Vascular Remodeling. Trans Am Clin Climatol Assoc. 2018;129:132–9.

60. Kim KK, Sheppard D, Chapman HA. TGF-beta1 Signaling and Tissue Fibrosis. Cold Spring Harb Perspect Biol. 2018;10(4).

61. Kim S, Cho S, Kim JH. CD1-mediated immune responses in mucosal tissues: molecular mechanisms underlying lipid antigen presentation system. Exp Mol Med. 2023;55(9):1858–71.

62. Wallace JM, Horgan GW, Bhattacharya S. Placental weight and efficiency in relation to maternal body mass index and the risk of pregnancy complications in women delivering singleton babies. Placenta. 2012;33(8):611–8.

63. Rosado-Yepez PI, Chavez-Corral DV, Reza-Lopez SA, Leal-Berumen I, Fierro-Murga R, Caballero-Cummings S, et al. Relation between pregestational obesity and characteristics of the placenta. J Matern Fetal Neonatal Med. 2020;33(20):3425–30.

64. Sureshchandra S, Doratt BM, True H, Mendoza N, Rincon M, Marshall NE, et al. Multimodal profiling of term human decidua demonstrates immune adaptations with pregravid obesity. Cell Rep. 2023;42(7):112769.

65. Malek R, Soufi S. Pulmonary Edema. StatPearls. Treasure Island (FL)2025.

66. Edwards Z, Annamaraju P. Physiology, Pulmonary Compliance. StatPearls. Treasure Island (FL)2025.

67. Sergienko NM, Bell JR, Weeks KL. Maternal obesity: influencing the heart right from the start. J Physiol. 2022;600(13):3007–8.

68. Ramos-Mondragon R, Galindo CA, Avila G. Role of TGF-beta on cardiac structural and electrical remodeling. Vasc Health Risk Manag. 2008;4(6):1289–300.

69. Kim M, Oh JK, Sakata S, Liang I, Park W, Hajjar RJ, et al. Role of resistin in cardiac contractility and hypertrophy. J Mol Cell Cardiol. 2008;45(2):270–80.

70. Bringhenti I, Moraes-Teixeira JA, Cunha MR, Ornellas F, Mandarim-de-Lacerda CA, Aguila MB. Maternal obesity during the preconception and early life periods alters pancreatic development in early and adult life in male mouse offspring. PLoS One. 2013;8(1):e55711.

71. Kahraman S, Dirice E, De Jesus DF, Hu J, Kulkarni RN. Maternal insulin resistance and transient hyperglycemia impact the metabolic and endocrine phenotypes of offspring. Am J Physiol Endocrinol Metab. 2014;307(10):E906–18.

72. Anastassova-Kristeva M. The origin and development of the immune system with a view to stem cell therapy. J Hematother Stem Cell Res. 2003;12(2):137–54.

73. Feltis BN, Wignarajah D, Zheng L, Ward C, Reid D, Harding R, et al. Increased vascular endothelial growth factor and receptors: relationship to angiogenesis in asthma. Am J Respir Crit Care Med. 2006;173(11):1201–7.

74. Stouch AN, McCoy AM, Greer RM, Lakhdari O, Yull FE, Blackwell TS, et al. IL-1beta and Inflammasome Activity Link Inflammation to Abnormal Fetal Airway Development. J Immunol. 2016;196(8):3411–20.

75. Yu VW, Scadden DT. Hematopoietic Stem Cell and Its Bone Marrow Niche. Curr Top Dev Biol. 2016;118:21–44.

76. Gao X, Xu C, Asada N, Frenette PS. The hematopoietic stem cell niche: from embryo to adult. Development. 2018;145(2).

77. Nakamura-Ishizu A, Ito K, Suda T. Hematopoietic Stem Cell Metabolism during Development and Aging. Dev Cell. 2020;54(2):239–55.

78. Wohnhaas CT, Bassler K, Watson CK, Shen Y, Leparc GG, Tilp C, et al. Monocyte-derived alveolar macrophages are key drivers of smoke-induced lung inflammation and tissue remodeling. Front Immunol. 2024;15:1325090.

79. Roche PA, Furuta K. The ins and outs of MHC class II-mediated antigen processing and presentation. Nat Rev Immunol. 2015;15(4):203–16.

80. Lampropoulou V, Sergushichev A, Bambouskova M, Nair S, Vincent EE, Loginicheva E, et al. Itaconate Links Inhibition of Succinate Dehydrogenase with Macrophage Metabolic Remodeling and Regulation of Inflammation. Cell Metab. 2016;24(1):158–66.

81. Wang Y, Branicky R, Noe A, Hekimi S. Superoxide dismutases: Dual roles in controlling ROS damage and regulating ROS signaling. J Cell Biol. 2018;217(6):1915–28.

82. Antonioli L, Pacher P, Vizi ES, Hasko G. CD39 and CD73 in immunity and inflammation. Trends Mol Med. 2013;19(6):355–67.

83. Deaglio S, Dwyer KM, Gao W, Friedman D, Usheva A, Erat A, et al. Adenosine generation catalyzed by CD39 and CD73 expressed on regulatory T cells mediates immune suppression. J Exp Med. 2007;204(6):1257–65.

84. Biernacka A, Dobaczewski M, Frangogiannis NG. TGF-beta signaling in fibrosis. Growth Factors. 2011;29(5):196–202.

85. Schroder M, Melum GR, Landsverk OJ, Bujko A, Yaqub S, Gran E, et al. CD1c-Expression by Monocytes - Implications for the Use of Commercial CD1c+ Dendritic Cell Isolation Kits. PLoS One. 2016;11(6):e0157387.

86. Osborne GA, Zhang L, Ma F, Gharaee-Kermani M, Turnier JL, Victory AN, et al. Dermatomyositis is characterized by JAK1-mediated monocyte-driven vasculopathy and inflammation. Sci Transl Med. 2025;17(830):eaea9007.

87. Chang CH, Pearce EL. Emerging concepts of T cell metabolism as a target of immunotherapy. Nat Immunol. 2016;17(4):364–8.

88. Yui MA, Rothenberg EV. Developmental gene networks: a triathlon on the course to T cell identity. Nat Rev Immunol. 2014;14(8):529–45.

89. Haks MC, Belkowski SM, Ciofani M, Rhodes M, Lefebvre JM, Trop S, et al. Low activation threshold as a mechanism for ligand-independent signaling in pre-T cells. J Immunol. 2003;170(6):2853–61.

90. Bray M, Alper MG. Lambda light chain predominance as a sign of emerging lymphoma. Am J Clin Pathol. 1983;80(4):526–8.

91. Arner P, Sahlqvist AS, Sinha I, Xu H, Yao X, Waterworth D, et al. The epigenetic signature of systemic insulin resistance in obese women. Diabetologia. 2016;59(11):2393–405.

92. Tellier J, Nutt SL. Plasma cells: The programming of an antibody-secreting machine. Eur J Immunol. 2019;49(1):30–7.

93. Nutt SL, Hodgkin PD, Tarlinton DM, Corcoran LM. The generation of antibody-secreting plasma cells. Nat Rev Immunol. 2015;15(3):160–71.

94. Elsner RA, Shlomchik MJ. Germinal Center and Extrafollicular B Cell Responses in Vaccination, Immunity, and Autoimmunity. Immunity. 2020;53(6):1136–50.

95. Oleinika K, Slisere B, Catalan D, Rosser EC. B cell contribution to immunometabolic dysfunction and impaired immune responses in obesity. Clin Exp Immunol. 2022;210(3):263–72.

