## Supplementary figures and images for "Spontaneous Pregravid Obesity Reshapes Fetal Immune Ontogeny in a Nonhuman Primate Model"

### Supp Figure 1

#### Supplemental Figure 1

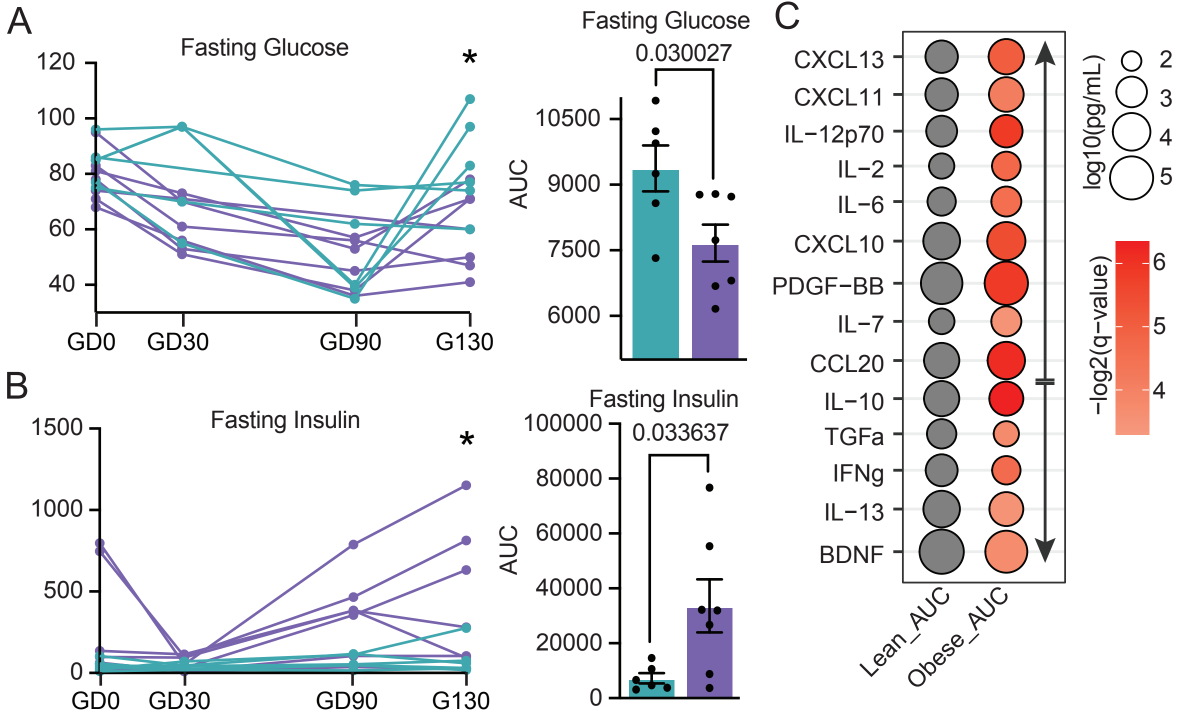

### Supp Figure 2

Supplemental Figure 2

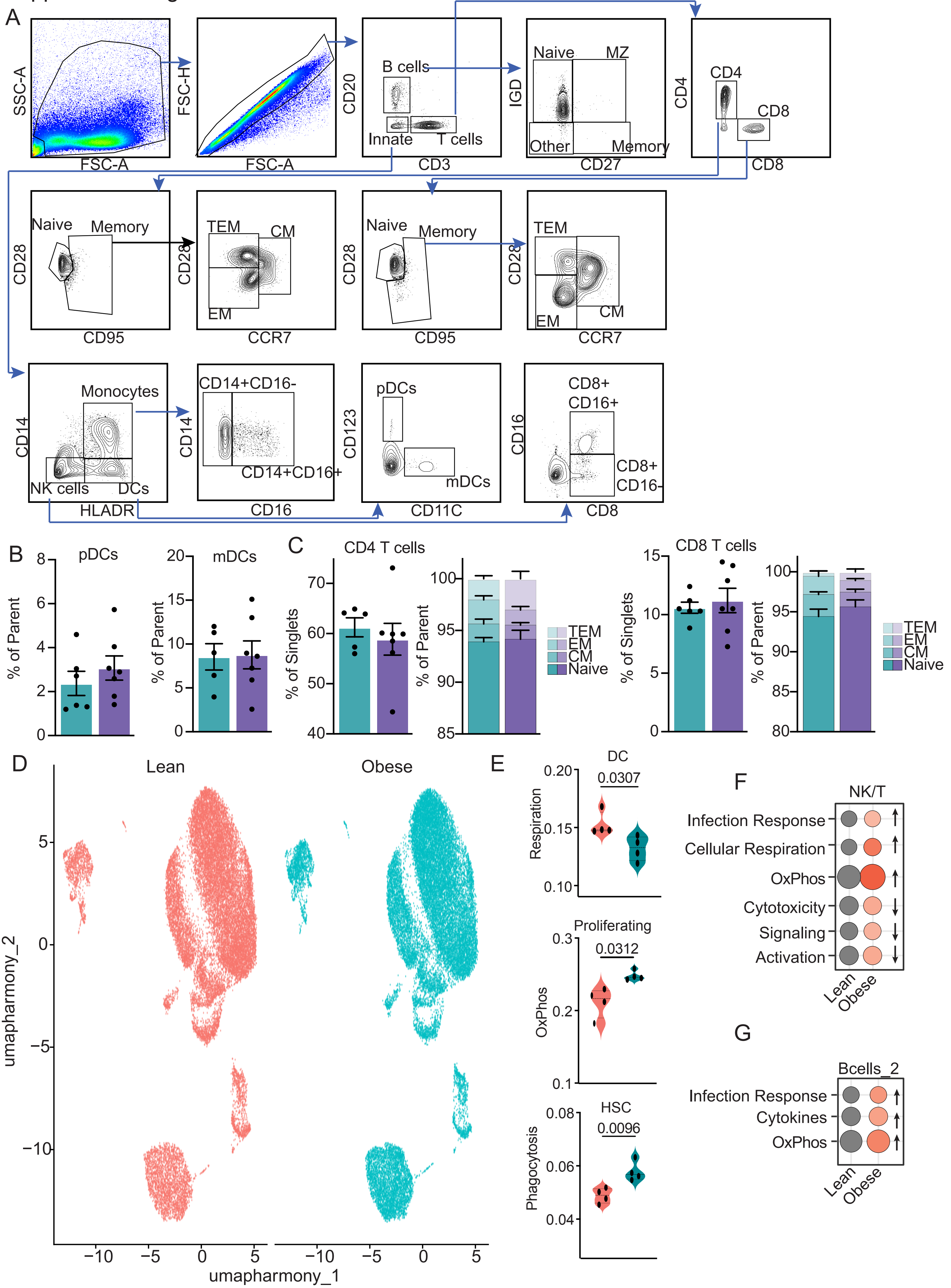

### Supp Figure 3

Supplemental Figure 3

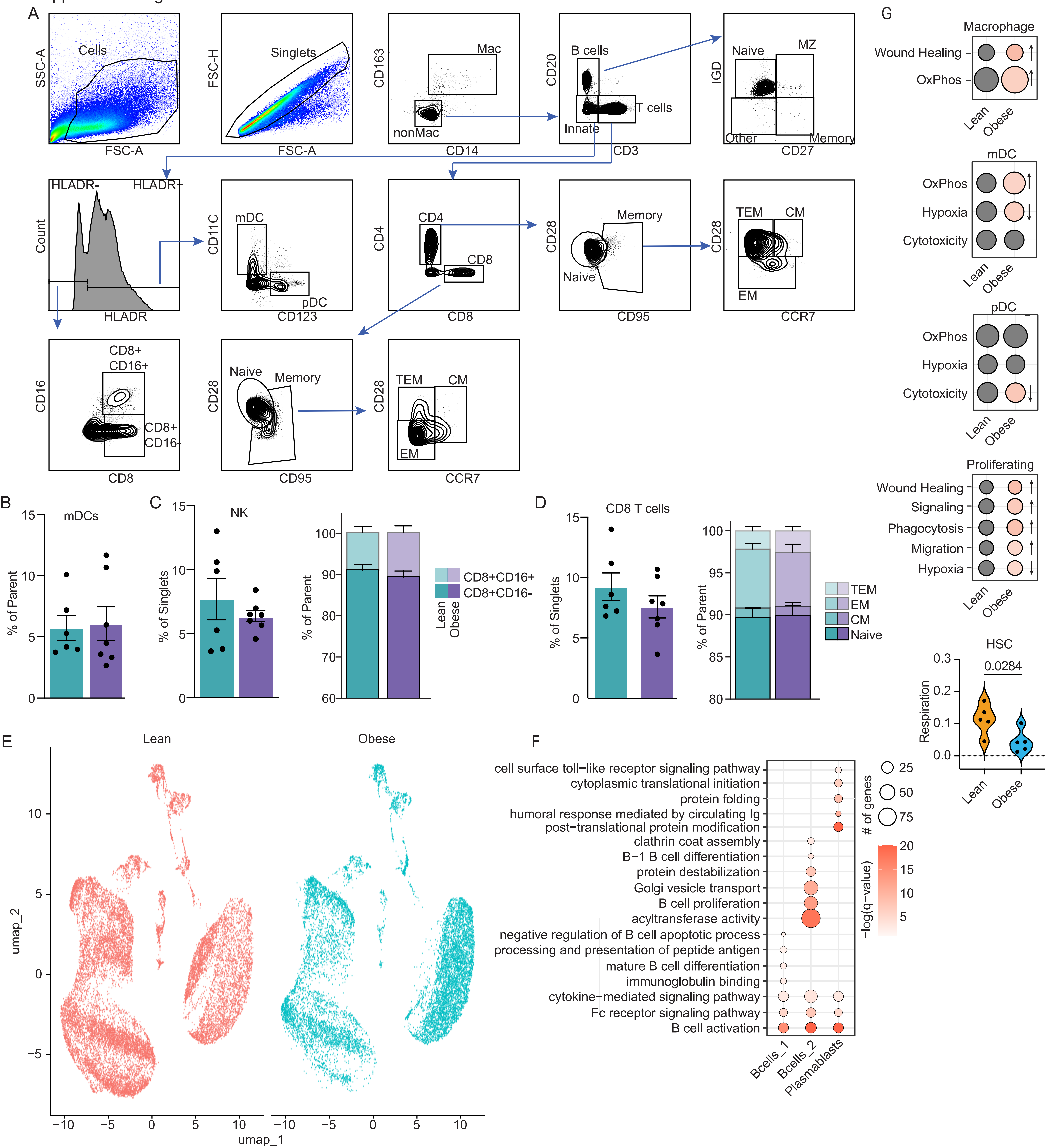

### Supp Figure 4

Supplemental Figure 4

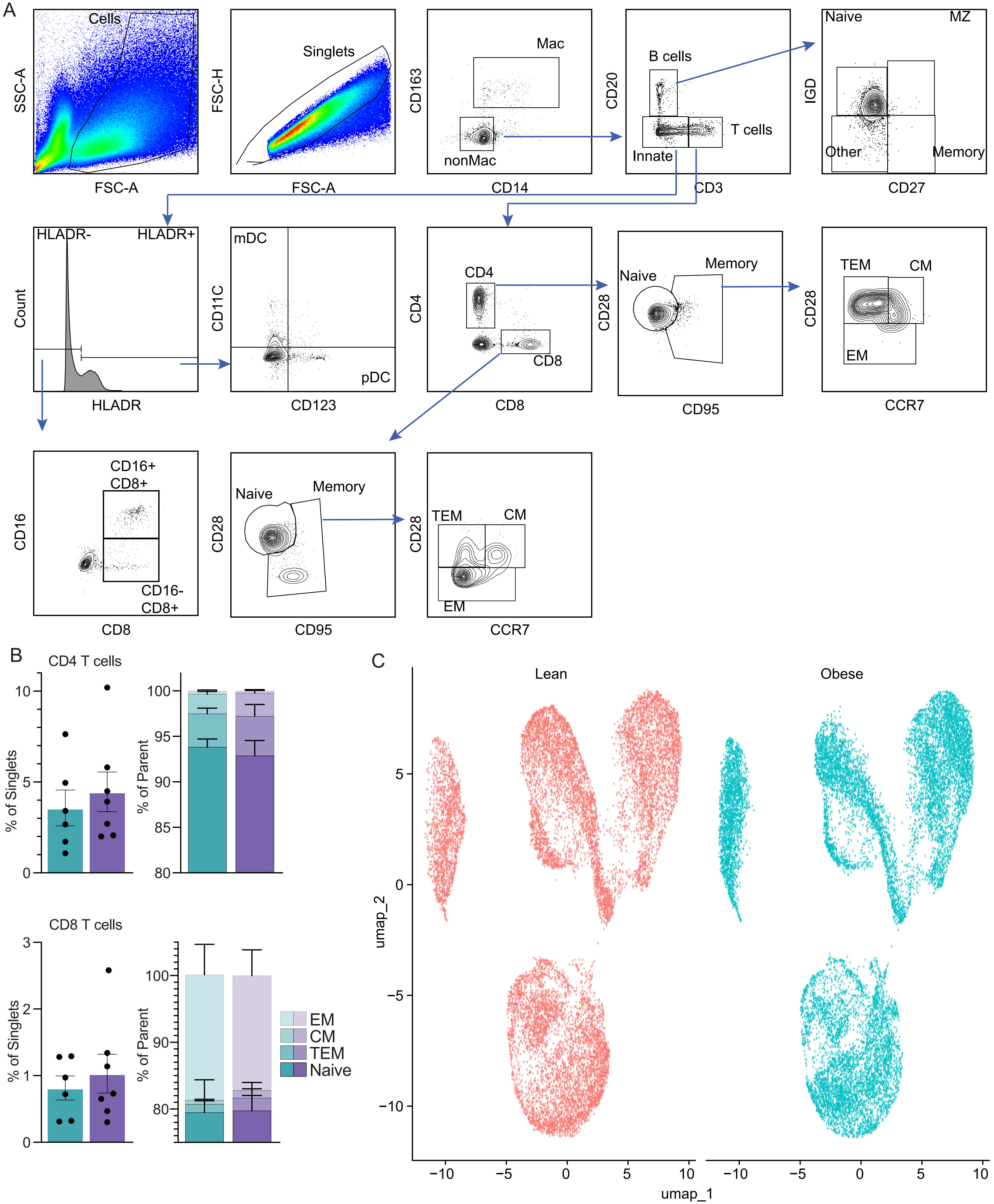
